# A scalable framework for high-throughput identification of functional origins of replication in non-model bacteria

**DOI:** 10.1101/2023.05.19.541510

**Authors:** Charlie Gilbert, Stephanie L. Brumwell, Alexander Crits-Christoph, Shinyoung Clair Kang, Zaira Martin-Moldes, Wajd Alsharif, Ariela Esmurria, Mary-Anne Nguyen, Henry H. Lee, Nili Ostrov

## Abstract

Microbial genetic manipulation requires access to engineerable plasmids that can be programmed to perturb genes, pathways and genomes. The extensive repertoire of plasmids available for model microbes, such as *Escherichia coli*, has facilitated fundamental biology studies and synthetic biology applications. However, the scarcity of plasmids for non-model microbes hinders efforts to broaden our biological knowledge and constrains the development of biotechnological solutions. In this study, we introduce a molecular toolkit and multiplexed screen to evaluate functional plasmids in non-model microbes. We constructed a collection of genetic parts consisting of 22 origins of replication (ORIs), 20 antibiotic selectable markers, and 30 molecular barcodes, which can be assembled combinatorially to create a library of plasmids trackable by next-generation DNA sequencing. We demonstrate our approach by delivering a pooled library of 22 ORIs to 12 bacterial species including extremophiles, electroactive bacteria and bioproduction strains. We report, for the first time, DNA delivery by conjugation and functional ORIs for *Halomonas alkaliphila, Halomonas neptunia,* and *Shewanella electrodiphila*. Furthermore, we expand the list of functional ORIs for *Duganella zoogloeoides*, *Pseudomonas alcaliphila*, *Shewanella oneidensis* and *Shewanella putrefaciens*. This screen provides a scalable high-throughput system to rapidly build and identify functional plasmids to establish genetic tractability in non-model microbes.

## Introduction

The availability of engineerable plasmids, replicating and programmable extrachromosomal pieces of DNA, opens up many opportunities for studying, perturbing, and programming an organism. Accordingly, plasmids are considered a fundamental tool for microbiology. Commonly used plasmids that replicate in *Escherichia coli*, however, are not always functional in other microbes. To facilitate the study and engineering of non-model microbes, it is thus desirable to establish a resource of plasmids that are functional in those organisms.

Most engineerable plasmids available today are derived from naturally-occurring plasmids, which can vary dramatically in size (∼1 kbp to >1 Mbp^1, 2^) and are not always found in a microbe of interest. Natural plasmids often carry additional genes with unknown functions. For recombinant DNA work in the laboratory, it is best to pare plasmids down to two key components: (1) a functional origin of replication (ORI) sequence responsible for DNA replication in the cell, and (2) a selectable marker, commonly encoding for antibiotic resistance which is used to maintain selection of the plasmid in the cell. Over time, excessive sequences and unnecessary genes have been empirically removed from natural plasmids to form compact engineerable plasmids which can be efficiently manipulated in the laboratory and transformed into cells^3–5^.

The molecular mechanisms governing an ORI’s host range are diverse and not always well understood. Plasmid origins replicate using a wide range of replication mechanisms which rely on and interact with different elements of the host’s cellular machinery^6, 7^. For instance, many ORIs encode proteins required for replication, and it is therefore vital that these genes are efficiently expressed in the host^8, 9^. ORIs may also contain DNA sequences or encode proteins that physically interact with host proteins, such as DNA polymerases or helicases; these interactions must remain efficient in non-native hosts for replication to occur^10, 11^. Overall, interactions between the host and plasmid must be preserved for ORIs to remain functional between species. As these interactions are difficult to predict, identifying functional ORIs for a particular microbe *a priori* is challenging, necessitating an empirical and often laborious approach.

To make the process of identifying functional engineerable plasmids in non-model microbes faster and more scalable, we developed a molecular toolkit containing multiple genetic parts, and an accompanying ORI-marker screen with a next-generation sequencing readout to efficiently explore large genetic design spaces (**Figure 1**). Our toolkit design allows for rapid exchange and combinatorial assembly of parts into barcoded ORI plasmid libraries. Our library of genetic parts comprises ORIs classified as broad host range (BHR), which are functional in multiple different species, genera, and even phyla, as well as ORIs classified as narrow host range (NHR) that are functional in just a handful of strains^1^. This library is compatible with multiple different antibiotic markers, making it widely applicable in any antibiotic-susceptible microbe. This collection of genetic parts is called the Possum (<u>P</u>lasmid <u>O</u>rigins and <u>S</u>electable Marker<u>s</u> for <u>U</u>ndomesticated <u>M</u>icrobes) Toolkit.

**Figure 1.**
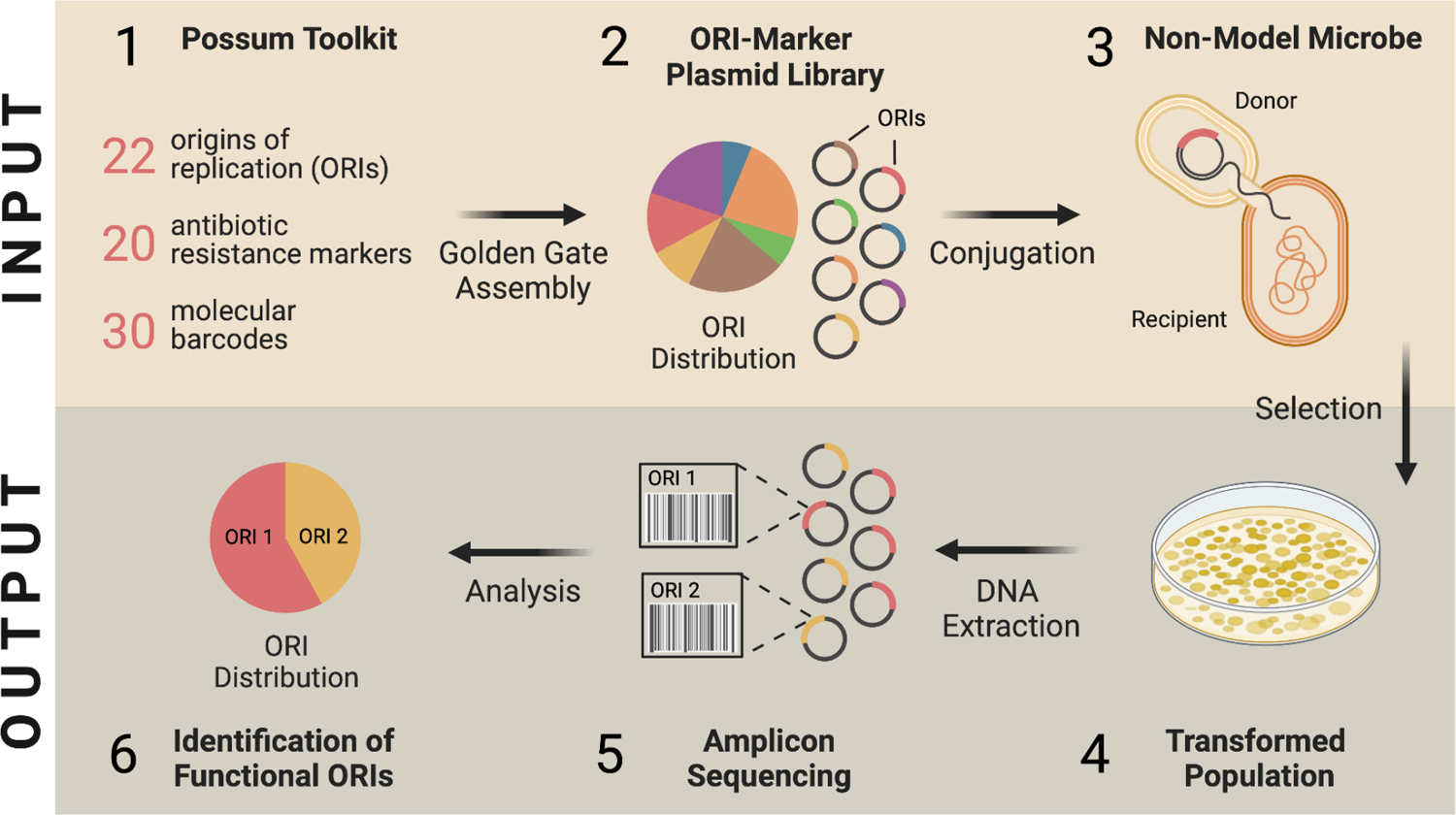
Overview of the ORI-marker screen. INPUT: (1) A pooled library of barcoded ORI-marker plasmids is built from genetic parts available in the Possum Toolkit via Golden Gate assembly. (2) The resulting plasmid library is delivered into an *E. coli* conjugative donor strain, and the distribution of ORI variants is determined by amplicon sequencing. (3) The library is then delivered to a target microbe through conjugation. OUTPUT: (4) Following conjugation, transformed cells are selected, pooled and then DNA is extracted to isolate plasmids. (5) NGS-based amplicon sequencing is performed on extracted DNA. (6) Data processing and statistical analyses detect functional ORIs from sequencing data. Created with BioRender.com.

## Results and Discussion

### ORI-marker library design

The Possum Toolkit provides a standardized, Golden-Gate compatible source of plasmid backbones and multiple genetic parts for non-model microbes. Our toolkit currently consists of 192 plasmids, including 22 BHR ORI variants and 20 antibiotic selectable markers (see **Data and Resource Availability**).

Our plasmid design allows cloning and handling of all constructs identically, regardless of whether the particular ORIs and antibiotic resistance genes being tested are also functional in *E. coli*. Specifically, all libraries in this study share the following plasmid backbone components: a conditional R6K ORI, a conjugative RP4 origin of transfer (*oriT*) and an *E. coli*-compatible chloramphenicol resistance gene (**Figure 2A**). The specific R6K ORI sequence we use (R6Kγ) lacks the native *pir* replication gene and therefore only functions in strains that have been engineered to express the *pir* gene^12, 13^. This means that the R6K ORI itself is extremely unlikely to work outside of a handful of *E. coli* cloning strains and will not interfere with testing of the ORI library. The *oriT* sequence was added to allow delivery of libraries by conjugation^14^. The chloramphenicol resistance gene is used to maintain the library plasmids in *E. coli* while cloning. We also added an mScarlet-I fluorescent reporter gene under the control of the constitutive bacterial promoter J23100 to help detect cells that are successfully transformed. It should be noted that mScarlet-I expression is likely to vary between microbes due to differences in promoter activity and plasmid copy number. Thus, the detection of mScarlet-I fluorescence alone in transformants is not sufficient to identify functional ORIs.

**Figure 2.**
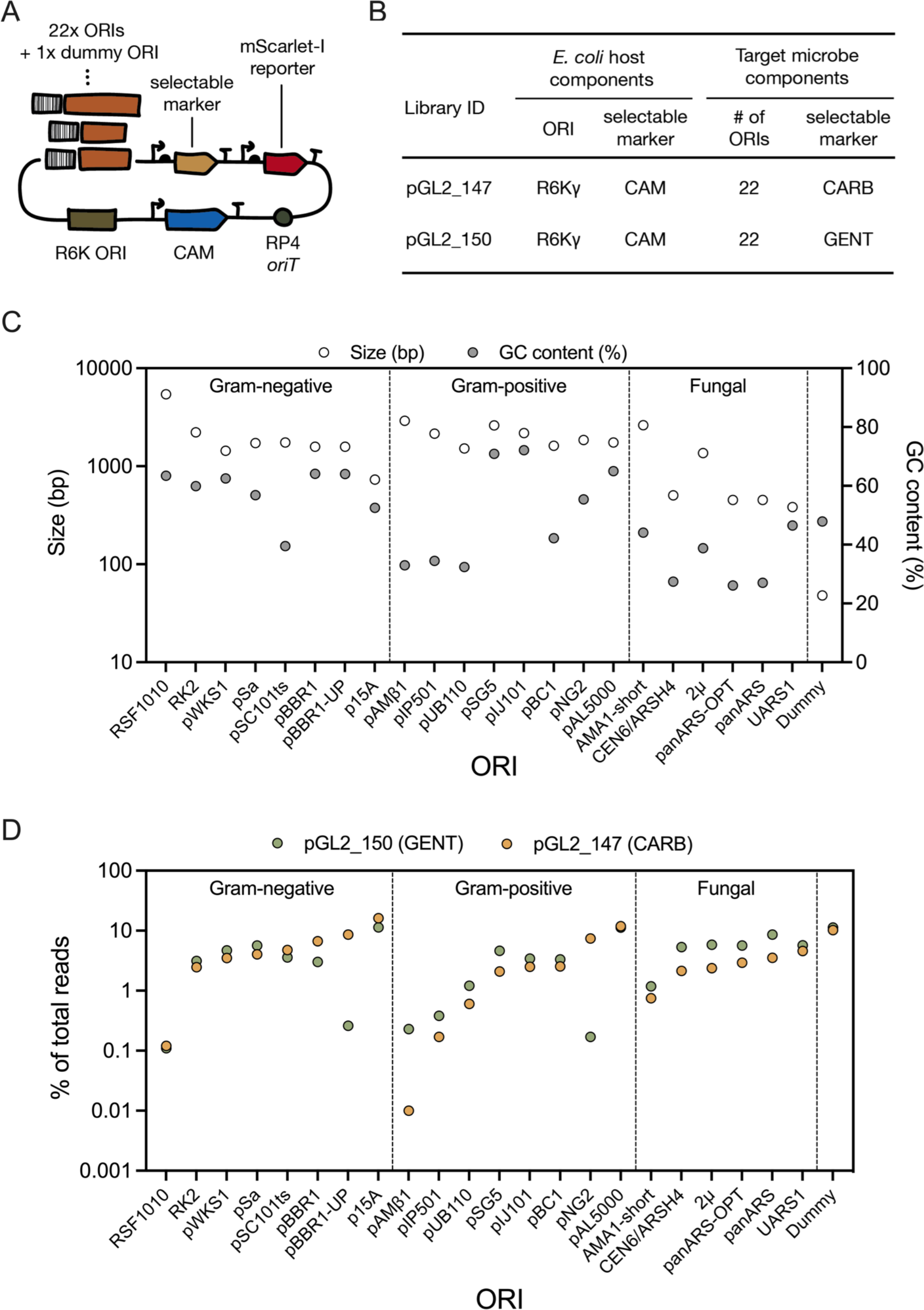
Plasmid library structure. (A) Schematic of the ORI-marker plasmid design. (B) Comparison of components in the pGL2_147 and pGL2_150 libraries. Both libraries were constructed with the same backbone, except for a carbenicllin (CARB) or gentamicin (GENT) antibiotic resistance marker. (C) Characterization of ORI size and GC content. ORIs are grouped based on the type of microbe they originated from: Gram-negative bacteria, Gram-positive bacteria or Fungi. The dummy ORI is a control sequence which should be non-functional in all microbes. (D) Distribution of ORIs in each input library as determined by amplicon sequencing of plasmids extracted from the respective conjugative donor strain. ORI distribution is represented as the percentage of total sequencing reads for each library.

In this work, we built two ORI-marker libraries: pGL2_147 with a carbenicillin (CARB) marker and pGL2_150 with a gentamicin (GENT) marker (**Figure 2B**). Each library contains the same set of 22 ORIs and a single non-functional ‘dummy’ ORI sequence that was added as an internal negative control (**Table 1, Supplementary Table 1**). Of the 22 ORI sequences tested, we classify 19 as having a broad host range and three as having a narrow host range. A total of 16 ORIs (68%) were originally isolated from prokaryotes and 6 ORIs (32%) from eukaryotes. The prokaryotic ORIs include eight that originate from Gram-positive species (pAL5000, pAMβ1, pIJ101, pIP501, pNG2, pSG5, pUB110, pBC1) and eight from Gram-negative species (p15A, pBBR1, pBBR1-UP, pSa, pSC101ts, pWKS1, RK2, RSF1010). All eukaryotic ORI sequences were originally isolated from fungi, either ascomycetes (2μ, AMA1 short, CEN6/ARSH4, panARS and panARS-opt) or basidiomycetes (UARS1). The average ORI size in our library is 1919 bp, ranging from 383 - 5435 bp (UARS1 and RSF1010, respectively). The average ORI GC-content is 48.8%, ranging from 26.1% - 72.1% (panARS-opt and pIJ101, respectively) (**Figure 2C**).

**Table 1.**
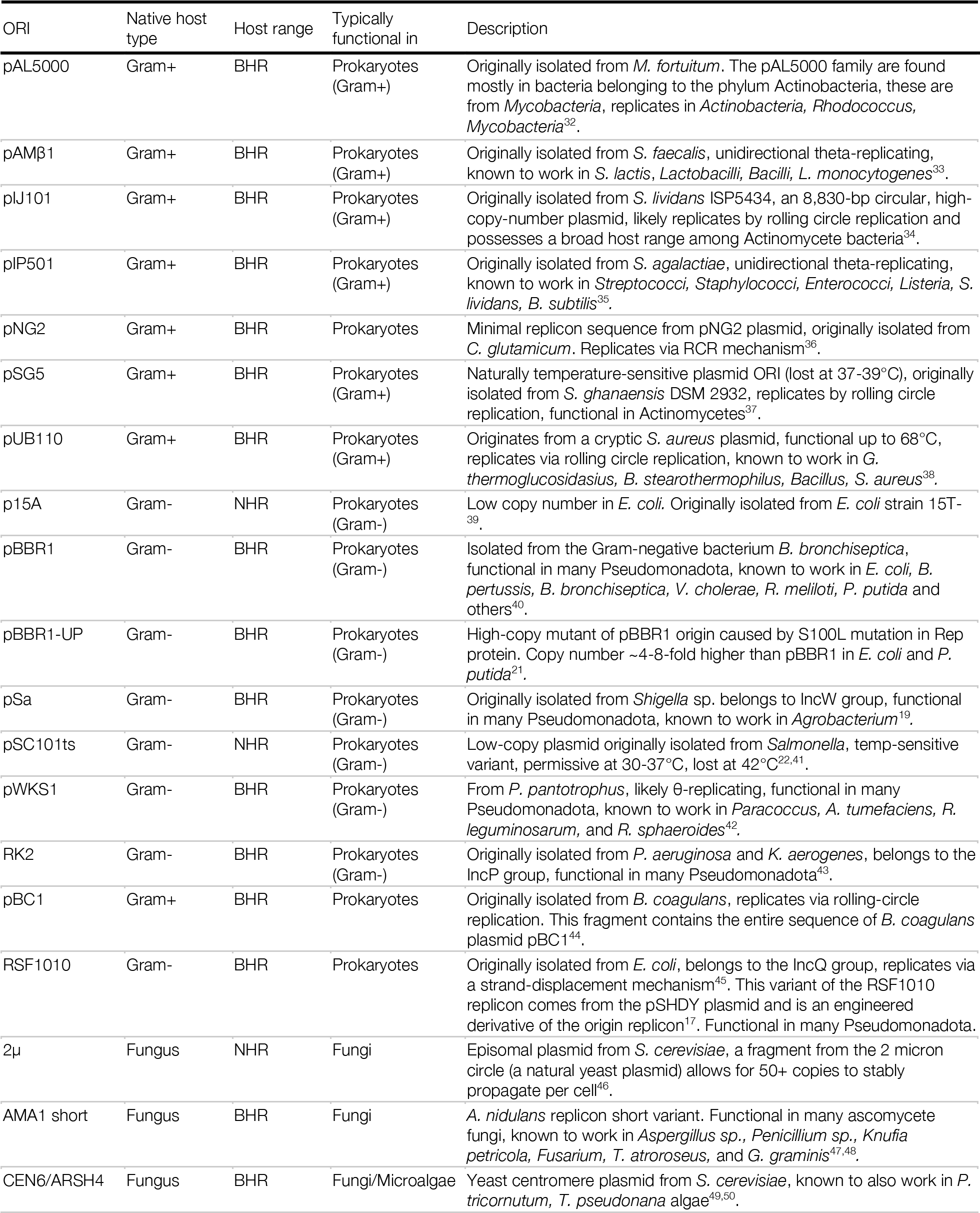
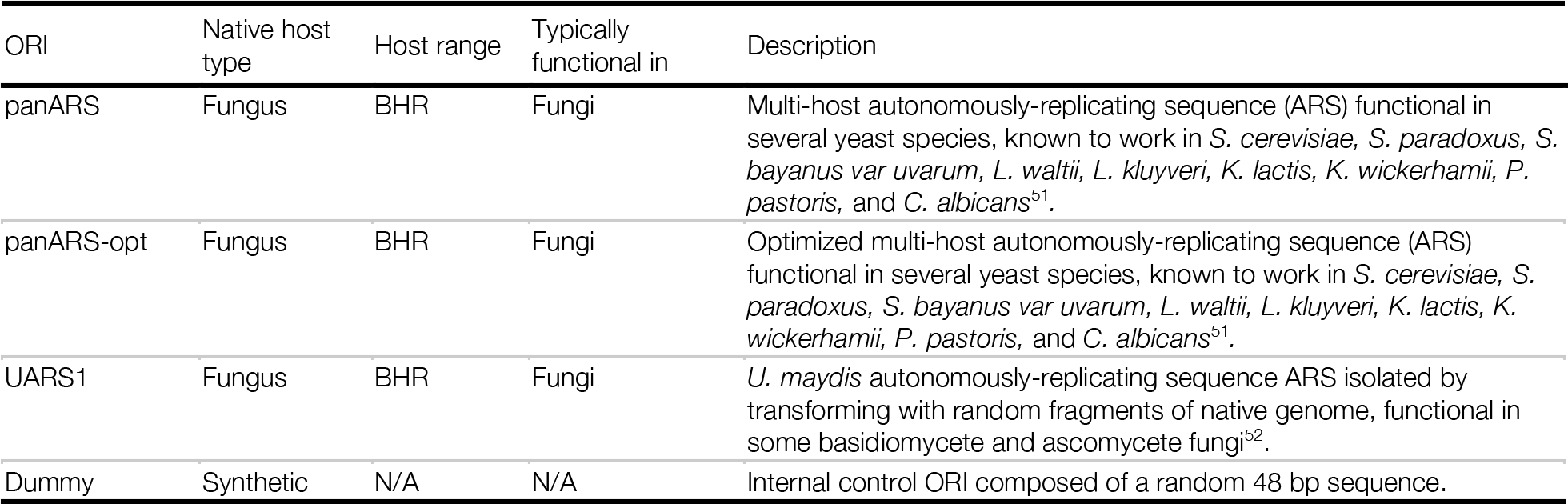
Origins of replication used in this study.

For conjugative delivery of plasmid libraries into target microbes, we selected the widely-used donor strain *E. coli* BW29427^15^. This strain contains the *pir* gene and RP4 conjugative machinery integrated into its genome and is a diaminopimelic acid (DAP) auxotroph allowing for easy counterselection and removal of donor cells after conjugation. We generated two donor strains carrying the “input library” by transforming *E. coli* BW29427 with pooled plasmid libraries pGL2_147 (CARB) or pGL2_150 (GENT).

We performed NGS on plasmid libraries extracted from the two donor strains to determine the relative abundance of ORIs in each input library (**Figure 2D**). All ORIs were represented in each input library. Most ORIs had similar abundance in both libraries (ranging from 1-10%), as may be expected based on the fact that all variants are maintained in the donor by the R6Kγ ORI. Some ORIs were consistently over or underrepresented in both pGL2_147 and pGL2_150 libraries, for example underrepresented ORIs RSF1010 (0.12% and 0.11%, respectively) and pAMβ1 (0.01% and 0.23%, respectively) and overrepresented ORI p15A (16.10% and 11.39%, respectively). Other ORIs exhibited substantial differences in abundance between libraries, such as pNG2 (7.42% vs 0.17%, respectively) and pBBR1-UP (8.61% vs 0.26%, respectively). Differences in individual ORI abundances could be caused by several factors such as ORI function in *E. coli*, GC-content, copy number or size.

### ORI-marker screen in non-model bacteria

We next set out to employ the ORI-marker screen to identify functional ORIs in 12 species of bacteria (**Table 2**). We used the following criteria to guide our choice of recipient bacteria: (1) under-studied species; (2) Gram-positive and Gram-negative (3 and 9 species, respectively), including nine *Pseudomonadota* (Proteobacteria), two *Bacillota* (Firmicutes) and one *Deinococcota*; (3) species with useful attributes *e.g.*, extremophiles that originate from unique environmental niches, including alkaliphiles, halophiles, psychrophiles, electroactive and radiation-resistant species; (4) organisms that were found to be compatible with *E. coli* growth conditions (*i.e.,* LB media, temperatures ranging from 20-37°C) to simplify co-culturing of donor and recipient and allow for a standardized assay across all microbes. We also included a common cloning strain of *E. coli* as a positive control recipient.

**Table 2.** Organisms tested in this ORI-marker screen.

| CVM ID | Strain | Strain ID | Phylum | Family | Gram | Attributes |
| --- | --- | --- | --- | --- | --- | --- |
| CVM022 | *Bacillus licheniformis* | ATCC 14580 | Bacillota | Bacillaceae | positive | bioproduction, probiotic |
| CVM028 | *Exiguobacterium aurantiacum* | ATCC BAA-333 | Bacillota | Exiguobacteraceae | positive | alkaliphile, halotolerant |
| CVM021 | *Deinococcus radiodurans* | ATCC 13939 | Deinococcota | Deinococcaceae | positive | radiation resistant, polyextremophile |
| CVM080 | *Escherichia coli* | Zymo 10B | Pseudomonadota | Enterobacteriaceae | negative | well-studied model |
| CVM032 | *Halomonas alkaliphila* | ATCC BAA-953 | Pseudomonadota | Halomonadaceae | negative | halophile |
| CVM027 | *Halomonas neptunia* | ATCC BAA-805 | Pseudomonadota | Halomonadaceae | negative | halophile, PHA production |
| CVM038 | *Duganella zoogloeoides* | ATCC 25935 | Pseudomonadota | Oxalobacteraceae | negative | EPS producer |
| CVM010 | *Pseudomonas alcaliphila* | ATCC BAA-571 | Pseudomonadota | Pseudomonadaceae | negative | psychrophile, alkaliphile, biosorption |
| CVM036 | *Shewanella electrodiphila* | ATCC BAA-2408 | Pseudomonadota | Shewanellaceae | negative | psychrophile, EPA producer, electroactive |
| CVM007 | *Shewanella oneidensis* | ATCC 700550 | Pseudomonadota | Shewanellaceae | negative | metal ion reduction, electroactive |
| CVM037 | *Shewanella putrefaciens* | ATCC 8071 | Pseudomonadota | Shewanellaceae | negative | metal ion reduction, electroactive |
| CVM029 | *Salinivibrio costicola* subsp. *alcaliphilus* | ATCC BAA-952 | Pseudomonadota | Vibrionales | negative | alkaliphile, halophile |

We tested library pGL2_150 (GENT) in all 12 recipients and library pGL2_147 (CARB) in five of the recipients, totaling 17 conjugation attempts. As a negative control, we performed conjugations using *E. coli* BW29427 lacking plasmid as a donor. Briefly, donor and recipient strains were grown separately at their respective optimal growth temperature and then mixed to a final donor-to-recipient ratio of 10:1 cells, as determined by OD_600_ (**Supplementary Table 2**). The mixture was plated on solid LB agar medium supplemented with DAP and incubated for 3 hours at a temperature optimal for the recipient strain. The conjugation mix was plated on selective and non-selective media using spot plating to determine conjugation frequency as well as plating on Petri dishes to maximize the number of transconjugants used for downstream NGS analysis (**Figure 3A**).

**Figure 3.**
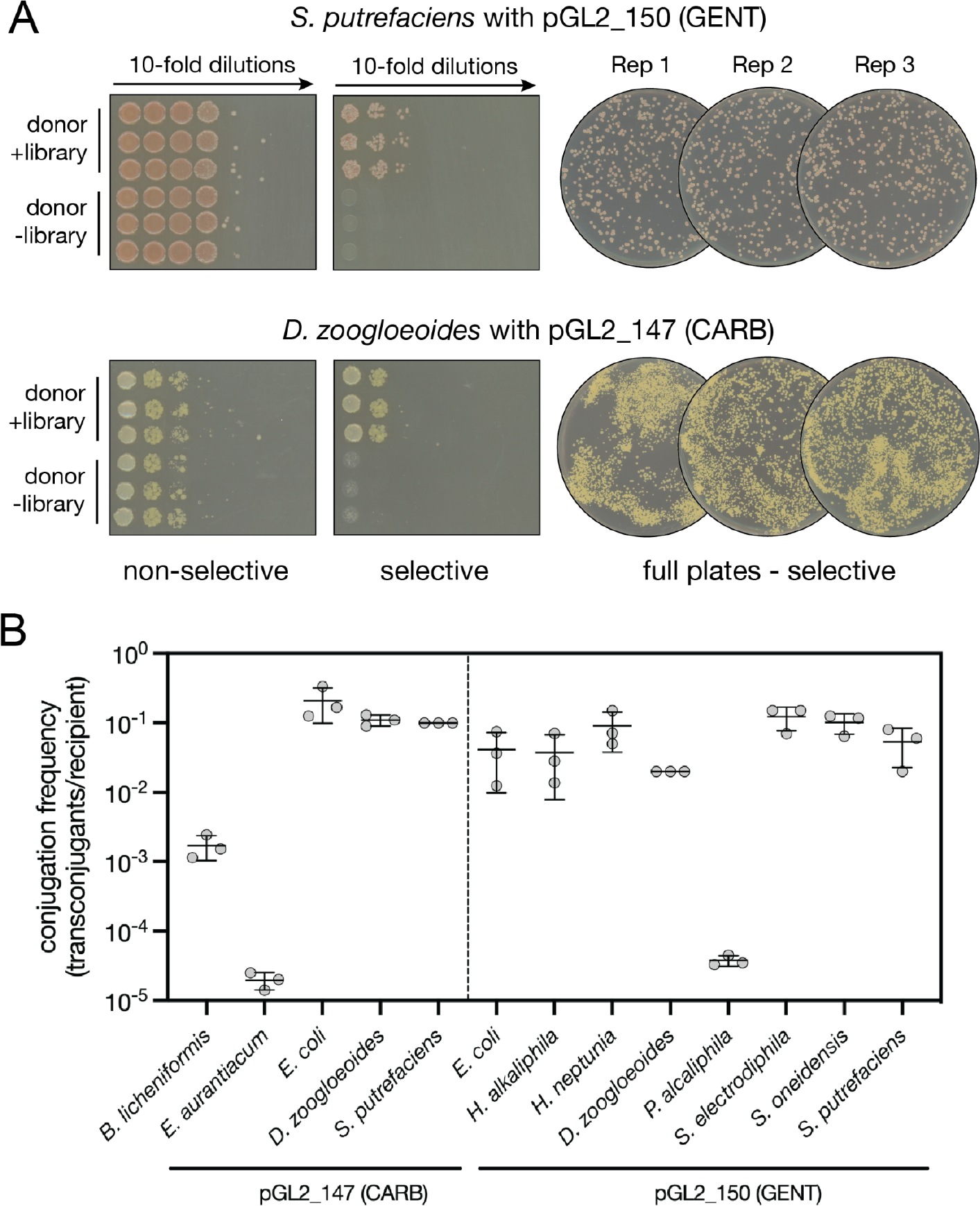
ORI-marker screen in non-model bacteria. (A) Representative images of recipient growth on spot plates and full plates following conjugation. *S. putrefaciens* (top) and *D. zoogloeoides* (bottom) following conjugation of the pGL2_150 (GENT) and pGL2_147 (CARB) libraries, respectively. 10-fold dilutions were spot plated on selective (LB-GENT^20^ or LB-CARB^100^) and non-selective (LB) plates (left), as well as selective media in Petri dishes (right). Conjugation was performed in triplicate (Rep 1-3) using an *E. coli* donor with (donor + library) or without the library (donor - library). (B) Conjugation frequency (transconjugants per recipient) for each recipient bacteria. Error bars represent standard deviation.

Overall, 13 of 17 (76%) conjugation attempts yielded colonies on selective media, corresponding to 10 of 12 recipient organisms (83%) (**Table 3**). We observed colonies for eight of 12 recipients with pGL2_150 (GENT) and all five recipients conjugated with pGL2_147 (CARB) (**Supplementary Table 3, Supplementary Figures 1 and 2**). Three recipient organisms yielded colonies with both libraries (*E. coli*, *Duganella zoogloeoides*, and *Shewanella putrefaciens*).

**Table 3.**
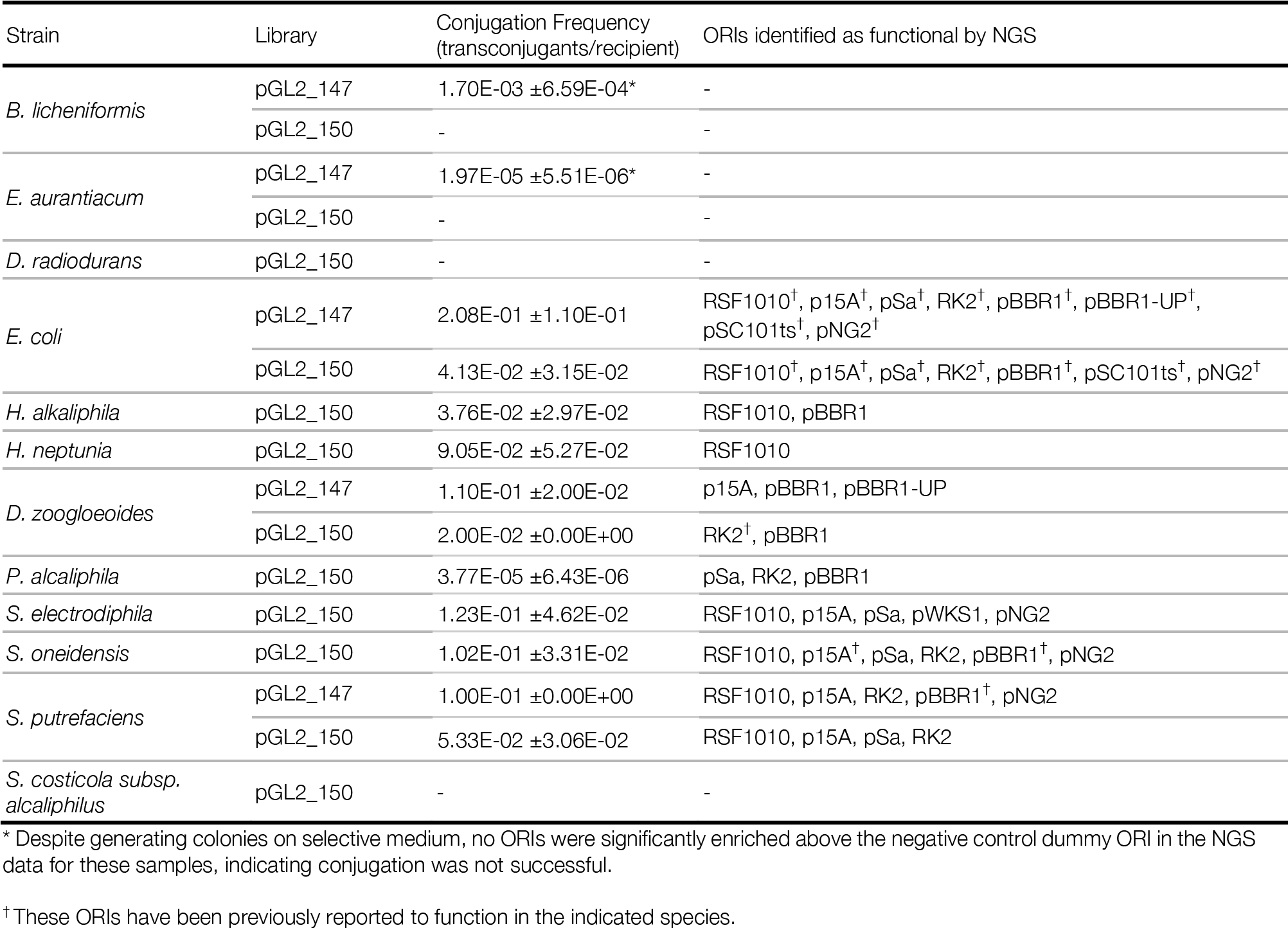
Summary of ORI-marker screen results.

No colonies were obtained in 4 of 17 conjugation attempts. There are a number of possible reasons for this. First, it is possible that our library does not contain an ORI that functions in these species. Second, as the efficiency of bacterial conjugation is known to vary between species, DNA delivery in these species may be too inefficient to generate transconjugants without optimization of our assay. Third, host restriction-modification systems or other foreign DNA defense mechanisms may have prevented plasmid replication. Finally, although all recipients are susceptible to the antibiotics used here, it is possible that the antibiotic resistance genes were not sufficiently expressed in these cases or that the antibiotic concentration used for selection was too high.

We calculated the conjugation frequency for each organism-library pair (**Figure 3B, Table 3, Supplementary Table 4**). Conjugation frequency of the pGL2_150 and pGL2_147 library from *E. coli* donors ranged from 10^-5^ to 10^-1^ transconjugants per recipient cell. Variations in conjugation frequency between recipients could be attributed to differences in functional ORIs (and therefore differences in the number of plasmids that can be delivered and replicated), standardized rather than optimized conjugation conditions, as well as recipient-specific conjugation dynamics; for example, the presence of active restriction-modification systems.

From the 13 samples that generated colonies on selective media, we harvested biomass for DNA extraction. Since conjugation protocols typically involve plating highly-concentrated cell suspensions, in some cases non-specific growth was observed for lower-dilution samples (**Supplementary Figures 1 and 2**). To account for this, colonies were only harvested where the growth of library recipient colonies on selective plates was significantly greater than the growth of control (no library) recipient cells.

### Identification of functional ORIs

To determine the abundance of functional ORIs in each recipient, we extracted DNA from transconjugants grown on selective media and performed NGS analysis. Notably, isolating DNA from non-model microbes presents species-specific difficulties that can impede efficient DNA extraction. For instance, *D. zoogloeoides* is a floc-forming, exopolysaccharide producer^16^ which tends to aggregate in liquid and on plates, making it difficult to resuspend and lyse the cells and interfering with DNA separation from magnetic beads. Despite these challenges, we were able to perform both plasmid and genomic DNA extraction for all species, resulting in average yields ranging from 4.2-170.3 ng/µL and 12.4-628.3 ng/µL, respectively (**Supplementary Table 5**).

We performed amplicon sequencing to identify functional ORIs in each recipient. Functional ORIs were defined as those that had amplicon counts significantly higher than the internal negative control “dummy” ORI (see **Targeted amplicon sequencing** Methods section). These analyses were designed to filter out false positive results, which could come from non-specific events such as random integration of the plasmid into genomic DNA, spontaneous mutants, or barcode amplification originating from the conjugation donor strain. Notably, the relative amplicon abundance observed for each ORI is influenced by several factors, including abundance in the input library, number of transconjugants obtained, transconjugant growth rate and plasmid copy number. Quantitative comparison between ORIs reported as functional in a given recipient should be treated with these factors in mind. Therefore, we use the rank order of ORI abundances as a high-level indicator of overall ORI functionality.

Overall, we were able to identify functional ORIs in eight recipient organisms (**Figure 4A**, **Table 3**, **Supplementary Figure 3**). Specifically, amplicon sequencing provided ORI results in 11 of 13 recipient-library pairs that yielded transconjugant colonies (eight with library pGL2_150 (GENT) and three with library pGL2_147 (CARB)). In the two remaining samples of *Exiguobacterium aurantiacum* and *Bacillus licheniformis* conjugated with library pGL2_147 (CARB), no ORIs were significantly enriched compared to the dummy ORI internal control, despite the fact that colonies were obtained on selective medium (**Supplementary Figure 2**). This indicates that the colonies obtained on selective media were able to survive through a non-specific process, such as random integration of the plasmid into chromosomal DNA or residual amounts of *E. coli* donor cells causing carbenicillin degradation. These results highlight the specificity of our NGS-based method which uses sequencing data, rather than colony count, to distinguish true transconjugants from false positives.

**Figure 4.**
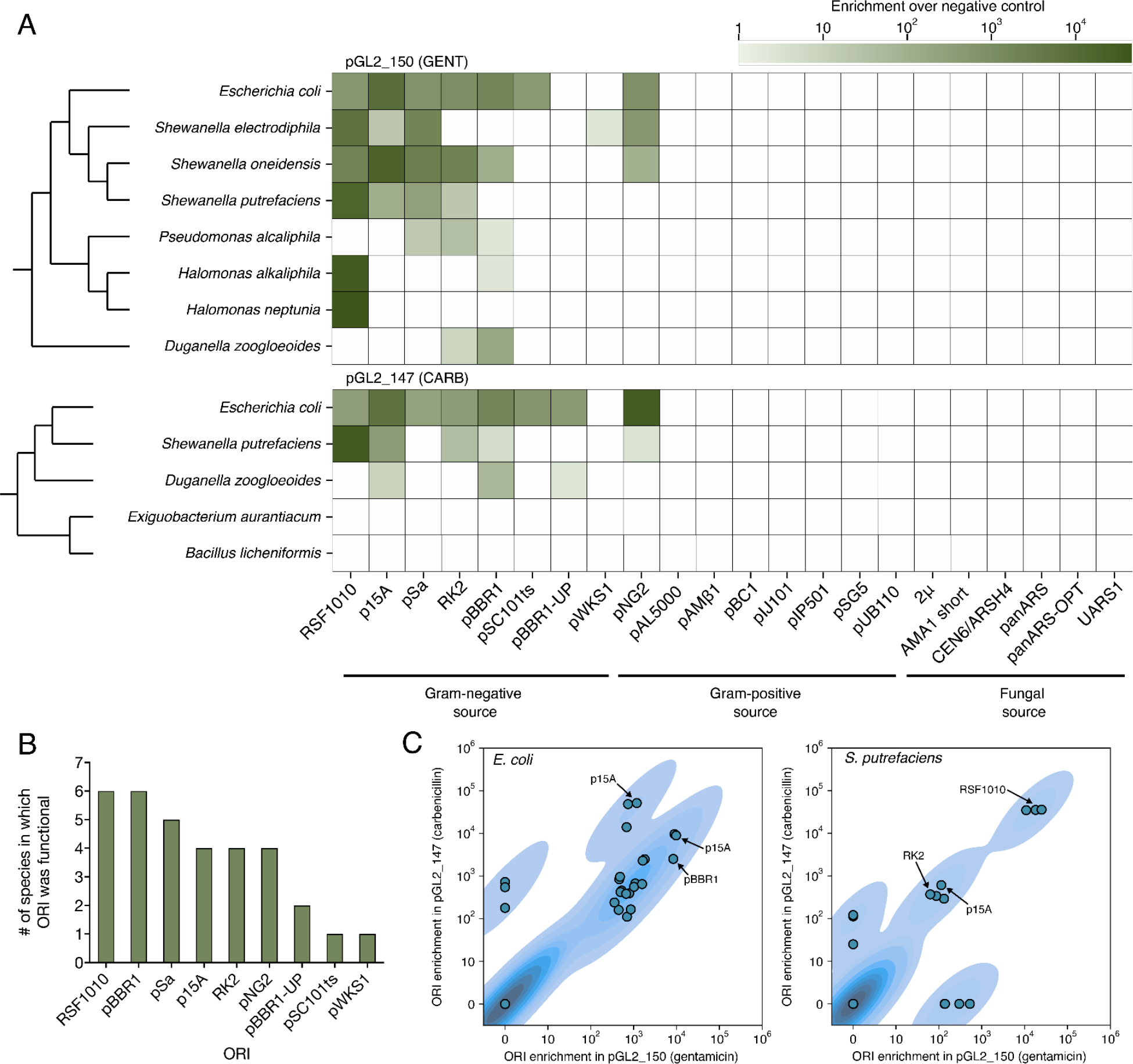
Summary of functional ORIs identified for 8 bacterial species. (A) Heat map of ORIs significantly enriched over non-functional control, as identified by amplicon sequencing in 8 bacteria. Organisms are shown with respective conjugated library pGL2_150 (GENT) (top) or pGL2_147 (CARB) (bottom). (p-adjusted < 0.01; see *Methods*). (B) Functional ORIs identified in this study and the respective number of bacteria for each. (C) Comparisons of ORI abundances as found in two strains which received both libraries pGL2_147 and pGL2_150.

We were encouraged to find that previously reported ORIs were recapitulated by our assay, including eight functional ORIs in *E. coli* (RSF1010^17^, p15A^18^, pSa^19^, RK2^14^, pBBR1^20^, pBBR1-UP^21^, pSC101ts^22^, pNG2^3^), as well as *D. zoogloeoides* (RK2^23^), *S. putrefaciens* (pBBR1^24^) and *Shewanella oneidensis* (p15A^25^, pBBR1^25^, pSC101^26^) (**Table 3**, **Supplementary Table 6**).

Importantly, we report here the first functional plasmids and ORIs for three non-model species, *H. alkaliphila, H. neptunia,* and *S. electrodiphila*, as well as previously unreported functional ORIs for *D. zoogloeoides*, *Pseudomonas alcaliphila*, *S. oneidensis* and *S. putrefaciens*. We successfully identified ORIs for all but one Gram-negative bacteria from the *Pseudomonadota* phylum, *Salinivibrio costicola* subsp. *alcaliphilus*. We observed two colonies on selective media for a single replicate of *S. costicola* with pGL2_150 (GENT), suggesting low efficiency of conjugation under the condition tested. For cases such as this, optimization of conjugation conditions (*e.g.*, donor-to-recipient ratio, media, duration) may increase the number of colonies obtained and allow functional ORI identification^27^.

We were unsuccessful in identifying functional ORIs in the three Gram-positive recipients tested in this work. One species from the Deinococcota phylum, *D. radiodurans*, yielded no colonies. Both species from the Bacillota phylum, *B. licheniformis and E. aurantiacum*, showed low-frequency conjugation compared to other Gram-negative recipients (0.0017 and 0.00002, respectively, corresponding to 47- and 4024-fold lower frequency compared with the average frequency in Gram-negative recipients) (**Figure 3B**). These putative transconjugants did not yield statistically significant sequencing abundances compared to the control origin to indicate the presence of functional ORIs and are therefore false positives. We hypothesize that the conjugation assay used in this work, which features a Gram-negative donor and a protocol largely optimized for Gram-negative recipients, is the major underlying cause of low-frequency conjugation. It is also possible that other selection markers or adjustment of the selection stringency would increase our ability to detect true transconjugants with functional ORIs. We are currently investigating improvements to expand the range of target microbes.

In total, nine of 22 ORIs in our library were found to be functional in one or more recipients (**Figure 4B**). In most recipients (7 of 8) we identified two or more functional ORIs, while only a single ORI (RSF1010) was identified for *H. neptunia*. ORIs RSF1010 and pBBR1 were the most common functional ORIs (detected in 6 of 8 recipients), while pSC101ts and pWKS1 were each found in only one recipient. This is despite the fact that RSF1010 was one of the least abundant library members in both input libraries (0.12% in pGL2_147 and 0.11% in pGL2_150), demonstrating the sensitivity of our screen and analysis pipeline, as even rare input library members can be amplified in compatible organisms. Our data also show the functionality of mesophilic ORIs in extremophiles that originate from different environmental niches.

We observed differences in the ORIs identified for the same recipient under different selective markers. Specifically, three organisms were attempted with both pGL2_147 (CARB) and pGL2_150 (GENT) libraries. In all of them, we observed at least one ORI identified with one selection marker but not the other (**Figure 4C, Table 3**). Specifically, in *E. coli* pBBR1-UP was only observed using pGL2_147; In *D. zoogloeoides*, pBBR1-UP and p15A were only observed using pGL2_147 while RK2 was only observed using pGL2_150; In *S. putrefaciens,* pBBR1 and pNG2 were only observed using pGL2_147 while pSa was only observed using pGL2_150. These differences may be caused by differences in ORI abundance in the input library (for example, pBBR1-UP abundance was 8.61% in pGL2_147 (CARB) vs 0.26% in pGL2_150 (GENT)), or due to differences in selection stringency between the selectable markers.

Transconjugants were further analyzed via whole plasmid sequencing to confirm the presence of ORIs reported by amplicon sequencing. While amplicon sequencing is expected to have greater sensitivity of detection, it may also be subject to technical issues that include but are not limited to amplification biases and amplicon contamination. In contrast, whole plasmid sequencing avoids these concerns, but lacks broad applicability due to differential lysis conditions for diverse species, and requires a more laborious sequencing library preparation protocol. We found a correlation of *r*=0.79 between the abundances of ORIs in amplicon and whole plasmid sequencing data, indicating a high degree of correspondence between the two methods (**Supplementary Figure 4**). Importantly, the top ORI identified by amplicon sequencing was confirmed to be present by whole plasmid sequencing for all 7 non-model strains and *E. coli* (**Supplementary Figures 3**). We note that some ORIs detected with lower read abundances were consistently reported by amplicon sequencing but could not be robustly detected with whole plasmid sequencing, which requires high sequencing depths. For this reason, we recommend using amplicon sequencing to maximize the sensitivity of functional ORI detection.

## Conclusion

Together, our Possum Toolkit and accompanying multiplexed sequencing screen provides a powerful resource to accelerate the identification of functional origins of replication in non-model microbes (**Table 4**). This pooled screen allowed us to quickly test 374 organism-plasmid combinations (*i.e.*, 22 ORIs each for 12 bacteria on plasmids with a gentamicin selection marker and for 5 bacteria on plasmids with a carbenicillin selection marker). With this assay, we were able to identify functional ORIs in 7 of 12 bacteria. We report successful DNA delivery by conjugation and functional ORIs in the halophilic bacteria *H. alkaliphila, H. neptunia*, and the electroactive bacterium *S. electrodiphila.* We also found additional functional ORIs in an exopolysaccharide-producer *D. zoogloeoides,* an alkaliphile *P. alcaliphila,* and electroactive bacteria *S. oneidensis* and *S. putrefaciens*. Importantly, our findings not only establish initial genetic tractability for these non-model microbes, but they also provide valuable negative data to further tease apart barriers and commonalities between microbial groups and inform development of impactful broad host range tools.

**Table 4.** Features of Cultivarium’s ORI-marker screen and Possum Toolkit.

| Feature | ORI-marker screen<br>(this publication) | Possum Toolkit<br>(genetic parts available through Addgene) |
| --- | --- | --- |
| Size | Each ORI-marker library contains 22 ORIs that can be screened simultaneously. | To our knowledge, represents the largest collection of Golden Gate-compatible plasmid ORIs for bacteria and fungi. |
| Broad host range | The 22x ORIs in our library are compatible with two fungal phyla and five bacterial phyla. |  |
| Selectable markers | We demonstrate the ORI-marker screen with two selectable markers: ampicillin/carbenicillin and gentamicin. | The Possum Toolkit contains 20 selectable markers that can be used to build plasmids for the ORI-marker screen. |
| Multiple DNA delivery modes | Our ORI-marker screen and Possum Toolkit parts are compatible with electroporation, bacterial conjugation and other standard DNA delivery modes. |  |
| Simple NGS readout | Uses a barcoded library approach, allowing screening of multiple ORIs in a single experiment. | Provides the parts and capability to add NGS barcodes to plasmid ORIs. |
| Resources available | We provide experimental and NGS analysis workflows to perform the ORI-marker screen. | Allows users to easily access our genetic parts library to clone their own plasmids. |
| Modularity and scalability | Plasmid libraries can be expanded and customized outside of the 22 ORIs used in this work. | The Possum Toolkit contains 192 genetic parts. This part collection will be expanded in the future by us and any potential users. |

These resources offer a rapid jumpstart for genetic manipulation in organisms for which no plasmid replication sequences are available. The current assay requires recipient organisms to be compatible with *E. coli* conjugation, which may be less applicable to some Gram-positive bacteria and fungi. To extend the utility of this approach, we are working to optimize conjugation conditions, expand the diversity of ORIs in the toolkit, and establish methods for high-throughput host-independent DNA delivery. As more functional ORIs are identified in diverse organisms, we expect the data sets from this high-throughput pooled ORI screen will reveal the rules governing compatibility of replication sequences and their hosts. Understanding these molecular mechanisms may allow the prediction of functional ORIs or enable *de novo* design of ORIs for non-model organisms.

## Materials and Methods

### Growth conditions

All strains used in this study were grown in LB broth or on LB 1.5% agar supplemented with diaminopimelic acid (DAP) 60 µg/mL, chloramphenicol (CAM) 34 µg/mL, carbenicillin (CARB) 100 µg/mL, and/or gentamicin (GENT) 20 µg/mL, when appropriate. Liquid cultures were grown with shaking at 225 RPM. *Escherichia coli* BW29427 (thrB1004 pro thi rpsL hsdS lacZΔM15 RP4-1360 Δ(araBAD)567 ΔdapA1341::[erm pir]) was grown at 37°C with DAP supplementation. All bacterial recipient strains and respective growth temperatures are summarized in **Supplementary Table 2**. The susceptibility of recipient strains to carbenicillin and gentamicin was established in solid and/or liquid cultures.

### ORI-marker library construction

The pGL2_147 (CARB) and pGL2_150 (GENT) input libraries were constructed by Golden Gate assembly using the Possum Toolkit, which adopts the Loop Assembly/uLoop Assembly Golden Gate standard^28, 29^. Barcoded ORIs, selectable marker genes, the mScarlet-I reporter gene and the *oriT* sequence were first assembled as Level 1 constructs from Level 0 parts. Next the selectable marker gene, mScarlet-I reporter gene and *oriT* sequence were assembled with a YFP dropout part sequence into a destination vector possessing the R6Kγ ORI and chloramphenicol resistance gene. This generated two Level 2 destination vectors, with either a CARB or GENT marker and a YFP dropout part occupying the assembly position which is filled by barcoded ORIs in the subsequent assembly step. In the final assembly step, 23x barcoded ORI Level 1 plasmids were combined in an equimolar pool and assembled into the YFP dropout part position. Finally, library transformants were pooled and plasmid DNA was extracted and transformed into the conjugation donor strain *E. coli* BW29427. All library variant sequences are hosted on GitHub at: https://github.com/cultivarium/ORI-marker-screen. Golden Gate assembly procedures were performed following standard Loop Assembly protocols^28^.

### Conjugation of ORI-marker libraries to recipient bacteria

#### Growth of recipient and donor strains

At least 2 days before conjugation, recipient strains were struck onto LB plates and incubated at their respective growth temperature (**Supplementary Table 2**). Microbes with longer doubling times may require a longer incubation period (*e.g.*, *Deinococcus radiodurans*). We used both liquid-and agar-based methods of growing recipient cultures prior to conjugation as indicated in **Supplementary Table 2**, as some non-model microbes have a long doubling time or grow slowly in LB broth. For liquid cultures: 1 day before conjugation, a single colony was inoculated into 50 mL of LB media. The cultures were grown overnight at their respective temperatures. For cultures on agar plates: recipients were used directly from the fresh streak plate.

##### Control donor

The day before conjugation, a single colony of *E. coli* BW29427 was inoculated into 5 mL of LB DAP media in a culture tube. In the morning, saturated overnight culture was diluted 1:50 into 50 mL of LB DAP media. This control culture was grown for 2 h at 37°C alongside the library donor. *Library donor:* Following ORI-marker library assembly and outgrowth, *E. coli* BW29427 donor cultures harboring pGL2_147 or pGL2_150 were frozen in 1 mL aliquots and stored at −80°C. On the day of conjugation, 1 aliquot per donor/recipient pairing was thawed at room temperature and added to 50 mL of LB DAP CAM media. Cultures were grown for 2 h at 37°C.

Cultures were adjusted to support a 10:1 donor-to-recipient ratio. For liquid cultures: The optical density (OD_600_) of the cultures was measured. Cultures were transferred into a 50-mL conical tube and pelleted by centrifugation at 4,000*g* at 4°C for 10 min. The supernatant was discarded. Donor pellets were washed once with 1 mL of 1X PBS to remove any residual antibiotic and centrifuged again. Donor pellets were resuspended to OD_600_ 100 and recipient pellets to OD_600_ 10 in LB media - creating a 10:1 ratio of donor:recipient for conjugation. For recipient cultures on agar plates: 500 µL of LB media was inoculated with enough cells from the fresh streak plate to reach OD_600_ of 10.

#### Conjugation, quantification, and biomass harvesting

50 µL of donor and 50 µL of recipient cells were mixed and spread directly on a conjugation plate (LB DAP) using a spreader until dry. Note: before conjugation, conjugation plates were dried in a biosafety cabinet for 1 hour. Each conjugation donor/recipient pairing was performed in triplicate, including control conjugations. Conjugation plates were incubated at the growth temperature of the recipient strain (see Table 3) for 3 hours, except for *S. electrodiphila* which was conjugated at 20°C rather than its optimal temperature of 15°C.

Conjugation plates were scraped with 1 mL of 1X PBS to resuspend cells. This mixture was then transferred to a 1.5 mL Eppendorf tube and 200 µL was aliquoted into the first column of a 96-well plate. Next, 10-fold serial dilutions were performed across an entire row of a 96-well plate for each sample (20 µL of sample into 180 µL of 1X PBS).

Spot plates were used to determine conjugation frequency. From each well of the 96-well dilution plate, 5 µL was spotted onto nonselective (LB) and selective (LB CARB or GEN) plates (in OmniTrays). Note: before spot plating, plates were dried in a biosafety cabinet for 1 hour. Spots were allowed to dry completely and then plates were incubated at the optimal temperature for recipient growth (as indicated in **Supplementary Table 2**) until colonies appeared. Colonies were counted manually. Conjugation frequency was calculated by dividing the number of colonies on selective spot plates by the number of colonies on nonselective spot plates (i.e., transconjugants/recipient), accounting for the dilution factor. Full plates were used to obtain biomass for NGS. 100 µL of 1-fold, 10-fold and 100-fold dilutions from the 96-well plate described above were plated onto selective media (in 100mm Petri dishes) using a spreader until dry. Plates were incubated at optimal growth temperature for the recipient until colonies appeared.

##### DNA extraction

For each replicate, the highest dilution plate with sufficient growth (aiming for 100+ colonies) was scraped with 1 mL of 1X PBS using a spreader to resuspend the transconjugants. In addition, three replicates of 10^-1^ and 10^-2^ dilution plates from the conjugation of pGL2_150 to *B. licheniformis* which contained no transconjugants were scraped and used as negative controls to assess background signal in amplicon sequencing. Colony counts for all plates from which colonies were harvested are listed in **Supplementary Table 7**. The cells were transferred to a 15-mL conical tube and the total volume was brought up to 2 mL with 1X PBS. When possible, we avoided collecting cells from the undiluted (10^0^) plate as this may have some background growth of the *E. coli* donor and the highest concentration of plasmid DNA from dead donor cells that could affect downstream NGS results. The OD_600_ was measured and cells were adjusted to a maximum OD_600_ of 10. The cells were split into two 1.5-mL Eppendorf tubes (1 mL each) and pelleted by centrifugation at room temperature at 12,000 RPM for 5 min. The supernatant was discarded. One of the two pellets was used for plasmid miniprep using the Qiagen Plasmid Miniprep Kit and the other for genomic DNA extraction using the MagMAX™ Viral/Pathogen Ultra Nucleic Acid Isolation Kit (standard volume 200-400 µL protocol, automated on KingFisher Flex) with the following modifications per sample: 2 µL of 100 ng/µL RNase was added to wash 1, 400 µL of binding solution and 40 µL of magnetic beads were used, and samples were eluted into 60 µL of elution buffer.

For some samples, the magnetic beads were not successfully removed from solution during the elution step (*B. licheniformis* pGL2_147 rep 2, *E. aurantiacum* pGL2_147 rep 2, *D. zoogloeoides* pGL2_147 all replicates and pGL2_150 rep 1 and 3, *S. electrodiphila* pGL2_150 all replicates). In these cases, an additional 200 µL of elution buffer was added, mixed by slow pipetting and transferred into a 1.5 mL Eppendorf tube. Samples were centrifuged at 12,000 RPM for 5 min to pellet the magnetic beads and the supernatant was collected (containing gDNA). The concentration of extracted DNA was determined using the Qubit™ 1X dsDNA High Sensitivity (HS) Assay Kit and adjusted to appropriate concentrations in elution buffer (10 mM Tris-Cl, pH 8.5). Genomic DNA was diluted to a maximum concentration of 10 ng/µL for amplicon sequencing. Plasmid DNA was adjusted to 1 ng/μL for whole plasmid sequencing.

##### Targeted amplicon sequencing of plasmid barcodes and analyses.

PCR 1 Mastermix was prepared by mixing 3.4 μL nuclease-free water, 5 μL of 2X Kapa Hifi HotStart Ready, 0.3 μL of PCR 1 Forward Primer (10 μM), and 0.3 μL PCR 1 Reverse Primer (10 μM). The PCR 1 primers (**Supplementary Table 8**) are designed to amplify the plasmid barcode sequence and to add stub sequences which bind to Illumina indexing primers in PCR 2.

For PCR 1 reaction, 9 μL of PCR 1 Mastermix was added to 1 μL of 10 ng/μL gDNA samples, then cycled in thermal cycler using the following optimized conditions: 95 °C for 3 min, 25 cycles of 20 sec denaturation at 98 °C, 15 sec annealing at 62 °C, and 30 sec extension at 72 °C, then cooling and hold at 4 °C. Concentrations of resulting amplicons were measured using Qubit fluorescence assay (Thermo Fisher), and each sample was diluted down to the concentration of 0.8-1 ng/μL.

PCR 2 adds unique dual indexed Illumina i5 and i7 primers to each of the samples via the complementarity of bases to the stub sequences added from PCR 1. PCR 2 Mastermix was prepared by mixing 4.5 μL of nuclease-free water and 7.5 μL of 2X Kapa Hifi HotStart Ready. 12 μL of PCR 2 Mastermix was added to each diluted sample, then 1 μL of i5xx (10 μM) and 1 μL of i7xx (10 μM) oligo primers were added. Following addition of the primers, the barcoded samples were cycled on the thermal cycler using the following optimized PCR 2 conditions: 72 °C for 3 min, 98 °C for 5 min, 10 cycles of 10 sec denaturation at 98 °C, 30 sec annealing at 62 °C, and 1 min of extension at 72 °C.

The barcoded samples were pooled together with equal volume, then purified with DNA Clean & Concentrator-5 (Zymo Research). The concentration of the purified, barcoded sample was measured using Qubit fluorescence assay, and the DNA fragment size of the amplicon was confirmed to have the expected size of 216 bp using Invitrogen E-Gel EX agarose gel. The final library was diluted to 4 nM using Resuspension Buffer (Illumina). The rest of the library preparation was completed following the Denature and Dilute Libraries Guide for MiniSeq System by Illumina, then loaded and sequenced on Illumina MiniSeq with a 300-cycle kit (2 x 150 bp paired-end run).

Amplicon sequencing reads (paired 2×150 bp) were first trimmed using BBDuk from the BBTools software package v39.01 (https://jgi.doe.gov/data-and-tools/software-tools/bbtools/) with the following settings: ktrim=r k=21 qtrim=r trimq=20 maq=20 minlen=50 entropy=0.3, and then merged with BBmerge using default settings. Trimmed and merged reads were matched against a database of barcode reference sequences associated with each ORI using vsearch^30^ with the setting -- search_exact; therefore, only perfect barcode matches were maintained. A median of 83% of sequenced reads passed trimming, merging, and barcode matching per sample.

The amplicon counts mapping to each barcoded ORI as a proportion of all reads were compared to the counts mapping to the non-functional “dummy” control origin within each sample using a two-sided z-test of proportions using the statsmodel package in Python v3.9. The resulting p-values were Bonferroni corrected, and ORIs significantly higher than the non-functional dummy’s proportion (p-adjusted<0.01) were retained. The same analysis was repeated on 6 negative control plates where donor DNA was plated at 10^-1^ and 10^-2^ dilutions but there were no successful conjugative colonies. The presence of DNA from the ORI pool library in these negative controls can be assumed to originate from the input library, and represents the abundance of the ORIs in the original library combined with any differential DNA decay that may occur during plating and incubation. Only ORIs with an enrichment over the non-functional dummy at least 5x greater than the maximum enrichment observed for any origin in the negative controls were reported as positive.

##### Whole plasmid sequencing and analyses

Tagmentation Mix was prepared by mixing 1.25 μL Tagment DNA Buffer and 0.25 μL Tagment DNA Enzyme (Illumina), then added to 1 μL of plasmid DNA at a concentration of 1 ng/μL for a total reaction volume of 2.5 μL. After gentle pipetting, the plasmids were incubated in the thermal cycler for 5 minutes at 55 °C, then allowed to cool to 4 °C. Samples were kept on ice until the next step.

PCR 2 (barcoding PCR) and the steps after were carried out exactly as the same as described previously for targeted sequencing of plasmid barcodes from gDNA, with the exception of using Illumina Nextera indexing primers - n5 and n7 - instead of i5 and i7 primers.

Pooled whole plasmid sequencing was also performed to further verify the presence and abundance of ORIs in transconjugants. Sequencing reads (paired 2×150 bp) were trimmed using BBDuk as described above. Reads were then mapped using Bowtie2 v2.4.4^31^ to a combined index of all plasmids belonging to the donor pool. Using Pysam 0.20, a custom script filtered reads that mapped to the unique ORI region of each plasmid, filtering to only reads mapped with at least 100 bp and 1 or fewer mismatches. Plasmid ORIs with at least 50% breadth of coverage of the origin region were called as present and the mean depth of coverage of the ORI region was reported.

## Data and Resource Availability

A major goal of this work is to provide the wider research community with a practical and user-friendly set of tools to expedite the genetic manipulation of non-model microbes of interest. Specifically, the library plasmids described in this work, as well as all genetic parts, plasmid backbones, and associated protocols, will be made available as part of the Possum Toolkit. Raw sequencing data, JupyterLab notebooks as well as scripts for the analysis, and results are hosted on GitHub at: https://github.com/cultivarium/ORI-marker-screen.

## Author Contributions

C.G., N.O. and H.H.L. conceived of and designed the project. Z.M.M., W.A. and A.E. established microbial culture conditions and antibiotic susceptibility. S.B. and C.G. performed library cloning and conjugation experiments. M.N. established automated genomic DNA extraction procedures. C.K. performed DNA sequencing workflows. A.C.C. analyzed the data. C.G., S.B., N.O., A.C.C., C.K. and H.H.L wrote the manuscript. All authors have read and approved the manuscript.

## Acknowledgements

We thank all members of the Cultivarium team for helpful discussions and technical advice throughout this project as well as helpful comments on this manuscript: Anna Douyon, Allison Lord, James Knight, Elise Ledieu-Dherbécourt, Michael Molla, Lilan Ling, Brianna Connolly, Julian Barros and Sierra Harken. Cultivarium acknowledges support from Schmidt Futures as a Convergent Research Focused Research Organization (FRO).

## Bibliography

1. Shintani, M., Sanchez, Z. K. & Kimbara, K. Genomics of microbial plasmids: classification and identification based on replication and transfer systems and host taxonomy. Front. Microbiol. 6, (2015).

2. Kothari, A. et al. Large Circular Plasmids from Groundwater Plasmidomes Span Multiple Incompatibility Groups and Are Enriched in Multimetal Resistance Genes. mBio 10, e02899–18 (2019).

3. Serwold-Davis, T. M., Groman, N. B. & Kao, C. C. Localization of an origin of replication in Corynebacterium diphtheriae broad host range plasmid pNG2 that also functions in Escherichia coli. FEMS Microbiol. Lett. 54, 119–123 (1990).

4. Cervantes-Rivera, R., Pedraza-López, F., Pérez-Segura, G. & Cevallos, M. A. The replication origin of a repABC plasmid. BMC Microbiol. 11, 158 (2011).

5. Tait, R. C., Close, T. J., Rodriguez, R. L. & Kado, C. I. Isolation of the origin of replication of the IncW-group plasmid pSa. Gene 20, 39–49 (1982).

6. Jain, A. & Srivastava, P. Broad host range plasmids. FEMS Microbiol. Lett. 348, 87–96 (2013).

7. del Solar, G., Giraldo, R., Ruiz-Echevarría, M. J., Espinosa, M. & Díaz-Orejas, R. Replication and Control of Circular Bacterial Plasmids. Microbiol. Mol. Biol. Rev. 62, 434–464 (1998).

8. Bartosik, D., Baj, J., Sochacka, M., Piechucka, E. & Wlodarczyk, M. Molecular characterization of functional modules of plasmid pWKS1 of Paracoccus pantotrophus DSM 11072. Microbiol. Read. Engl. 148, 2847–2856 (2002).

9. Brantl, S., Behnke, D. & Alonso, J. C. Molecular analysis of the replication region of the conjugative Streptococcus agalactiae plasmid pIP501 in Bacillus subtilis. Comparison with plasmids pAM beta 1 and pSM19035. Nucleic Acids Res. 18, 4783–4790 (1990).

10. Masai, H. & Arai, K. RepA and DnaA proteins are required for initiation of R1 plasmid replication in vitro and interact with the oriR sequence. Proc. Natl. Acad. Sci. U. S. A. 84, 4781–4785 (1987).

11. Ratnakar, P. V., Mohanty, B. K., Lobert, M. & Bastia, D. The replication initiator protein pi of the plasmid R6K specifically interacts with the host-encoded helicase DnaB. Proc. Natl. Acad. Sci. 93, 5522–5526 (1996).

12. Kolter, R., Inuzuka, M. & Helinski, D. R. Trans-complementation-dependent replication of a low molecular weight origin fragment from plasmid R6K. Cell 15, 1199–1208 (1978).

13. Stalker, D. M., Kolter, R. & Helinski, D. R. Nucleotide sequence of the region of an origin of replication of the antibiotic resistance plasmid R6K. Proc. Natl. Acad. Sci. U. S. A. 76, 1150– 1154 (1979).

14. Fürste, J. P., Pansegrau, W., Ziegelin, G., Kröger, M. & Lanka, E. Conjugative transfer of promiscuous IncP plasmids: interaction of plasmid-encoded products with the transfer origin. Proc. Natl. Acad. Sci. U. S. A. 86, 1771–1775 (1989).

15. Lee, H. H. et al. Functional genomics of the rapidly replicating bacterium Vibrio natriegens by CRISPRi. Nat. Microbiol. 4, 1105–1113 (2019).

16. Lee, S., Kang, S., Kwon, C. & Jung, S. Zooglan, an extracellular acidic polysaccharide isolated from Zoogloea ramigera 115 as a novel catalytic carbohydrate for methanolysis. Carbohydr. Polym. 64, 350–354 (2006).

17. Behle, A. & Axmann, I. M. pSHDY: A New Tool for Genetic Engineering of Cyanobacteria. In Methods in molecular biology (Clifton, N.J.) vol. 2379 67–79 (Methods Mol Biol, 2022).

18. Selzer, G., Som, T., Itoh, T. & Tomizawa, J. The origin of replication of plasmid p15A and comparative studies on the nucleotide sequences around the origin of related plasmids. Cell 32, 119–129 (1983).

19. Watanabe, T., Furuse, C. & Sakaizumi, S. Transduction of various R factors by phage P1 in Escherichia coli and by phage P22 in Salmonella typhimurium. J. Bacteriol. 96, 1791–1795 (1968).

20. Kovach, M. E. et al. Four new derivatives of the broad-host-range cloning vector pBBR1MCS, carrying different antibiotic-resistance cassettes. Gene 166, 175–176 (1995).

21. Tao, L., Jackson, R. E. & Cheng, Q. Directed evolution of copy number of a broad host range plasmid for metabolic engineering. Metab. Eng. 7, 10–17 (2005).

22. Armstrong, K. a et al. A 37 × 10(3) molecular weight plasmid-encoded protein is required for replication and copy number control in the plasmid pSC101 and its temperature-sensitive derivative pHS1. J. Mol. Biol. 175, 331–48 (1984).

23. Easson, D. D. A recombinant DNA approach to the design and synthesis of novel polysaccharides. (Massachusetts Institute of Technology, 1987).

24. Wei, H. et al. Functional roles of CymA and NapC in reduction of nitrate and nitrite by Shewanella putrefaciens W3-18-1. Microbiology 162, 930–941 (2016).

25. Towards enabling engineered microbial-electronic systems: RK2-based conjugal transfer system for Shewanella synthetic biology - Hajimorad - 2016 - Electronics Letters - Wiley Online Library. https://ietresearch.onlinelibrary.wiley.com/doi/full/10.1049/el.2015.3226.

26. Cao, Y. et al. A Synthetic Plasmid Toolkit for Shewanella oneidensis MR-1. Front. Microbiol. 10, (2019).

27. Vargas, C., Coronado, M. J., Ventosa, A. & Nieto, J. J. Host Range, Stability and Compatibility of Broad Host-Range-Plasmids and a Shuttle Vector in Moderately Halophilic Bacteria. Evidence of Intrageneric and Intergeneric Conjugation in Moderate Halophiles. Syst. Appl. Microbiol. 20, 173–181 (1997).

28. Pollak, B. et al. Loop assembly: a simple and open system for recursive fabrication of DNA circuits. New Phytol. 222, 628–640 (2019).

29. Pollak, B. et al. Universal loop assembly: Open, efficient and cross-kingdom DNA fabrication. Synth. Biol. 5, 1 (2019).

30. Rognes, T., Flouri, T., Nichols, B., Quince, C. & Mahé, F. VSEARCH: a versatile open source tool for metagenomics. PeerJ 4, e2584 (2016).

31. Langmead, B. & Salzberg, S. L. Fast gapped-read alignment with Bowtie 2. Nat. Methods 9, 357–359 (2012).

32. Rauzier, J., Moniz-Pereira, J. & Gicquel-Sanzey, B. Complete nucleotide sequence of pAL5000, a plasmid from Mycobacterium fortuitum. Gene 71, 315–321 (1988).

33. Bruand, C., Le Chatelier, E., Ehrlich, S. D. & Jannière, L. A fourth class of theta-replicating plasmids: the pAM beta 1 family from gram-positive bacteria. Proc. Natl. Acad. Sci. U. S. A. 90, 11668–11672 (1993).

34. Kieser, T., Hopwood, D. A., Wright, H. M. & Thompson, C. J. pIJ101, a multi-copy broad host-range Streptomyces plasmid: functional analysis and development of DNA cloning vectors. Mol. Gen. Genet. MGG 185, 223–228 (1982).

35. Horodniceanu, T., Bouanchaud, D. H., Bieth, G. & Chabbert, Y. A. R Plasmids in Streptococcus agalactiae (Group B). Antimicrob. Agents Chemother. 10, 795–801 (1976).

36. Schiller, J., Groman, N. & Coyle, M. Plasmids in Corynebacterium diphtheriae and diphtheroids mediating erythromycin resistance. Antimicrob. Agents Chemother. 18, 814–821 (1980).

37. Wohlleben, W. D., Schulte, A., Pühler, A. P. D. & Muth, G. Streptomycetes plasmid pSG5, process for its preparation and its use. (1985).

38. Gryczan, T. J., Contente, S. & Dubnau, D. Characterization of Staphylococcus aureus plasmids introduced by transformation into Bacillus subtilis. J. Bacteriol. 134, 318–329 (1978).

39. Cozzarelli, N. R., Kelly, R. B. & Kornberg, A. A minute circular DNA from Escherichia coli 15. Proc. Natl. Acad. Sci. 60, 992–999 (1968).

40. Antoine, R. & Locht, C. Isolation and molecular characterization of a novel broad-host-range plasmid from Bordetella bronchiseptica with sequence similarities to plasmids from gram-positive organisms. Mol. Microbiol. 6, 1785–1799 (1992).

41. Cohen, S. N. & Chang, A. C. Recircularization and autonomous replication of a sheared R-factor DNA segment in Escherichia coli transformants. Proc. Natl. Acad. Sci. U. S. A. 70, 1293–1297 (1973).

42. Baj, J., Piechucka, E., Bartosik, D. & Włodarczyk, M. Plasmid occurrence and diversity in the genus Paracoccus. Acta Microbiol. Pol. 49, 265–270 (2000).

43. Pansegrau, W. et al. Complete nucleotide sequence of Birmingham IncP alpha plasmids. Compilation and comparative analysis. J. Mol. Biol. 239, 623–663 (1994).

44. De Rossi, E., Brigidi, P., Riccardi, G. & Matteuzzi, D. Plasmid screening in thermophilicBacillus: Physical characterization and molecular cloning. Curr. Microbiol. 19, 13–19 (1989).

45. Guerry, P., van Embden, J. & Falkow, S. Molecular nature of two nonconjugative plasmids carrying drug resistance genes. J. Bacteriol. 117, 619–630 (1974).

46. Chan, K.-M., Liu, Y.-T., Ma, C.-H., Jayaram, M. & Sau, S. The 2 micron plasmid of Saccharomyces cerevisiae: A miniaturized selfish genome with optimized functional competence. Plasmid 70, 2–17 (2013).

47. Gems, D., Johnstone, I. L. & Clutterbuck, A. J. An autonomously replicating plasmid transforms Aspergillus nidulans at high frequency. Gene 98, 61–67 (1991).

48. Mózsik, L. et al. Modular Synthetic Biology Toolkit for Filamentous Fungi. ACS Synth. Biol. acssynbio.1c00260 (2021) doi:10.1021/ACSSYNBIO.1C00260.

49. Newlon, C. S. & Theis, J. F. The structure and function of yeast ARS elements. Curr. Opin. Genet. Dev. 3, 752–758 (1993).

50. Cottarel, G., Shero, J. H., Hieter, P. & Hegemann, J. H. A 125-base-pair CEN6 DNA fragment is sufficient for complete meiotic and mitotic centromere functions in Saccharomyces cerevisiae. Mol. Cell. Biol. 9, 3342–3349 (1989).

51. Liachko, I. & Dunham, M. J. An autonomously replicating sequence for use in a wide range of budding yeasts. FEMS Yeast Res. 14, 364–367 (2014).

52. Tsukuda, T., Carleton, S., Fotheringham, S. & Holloman, W. K. Isolation and characterization of an autonomously replicating sequence from Ustilago maydis. Mol. Cell. Biol. 8, 3703–3709 (1988).

