## Supplementary Materials for "A scalable framework for high-throughput identification of functional origins of replication in non-model bacteria"

**Supplementary Figures** **2**
Supplementary Figure 1. Conjugation plate images - Library pGL2_150. 3
Supplementary Figure 2. Conjugation plate images - Library pGL2_147 4
Supplementary Figure 3. Relative ORI abundances in each recipient bacteria. 9
Supplementary Figure 4. Correlation between amplicon and whole plasmid sequencing. 9
**Supplementary Tables** **10**
Supplementary Table 1. Nucleotide sequences of the ORIs used in this study. 10
Supplementary Table 2. Conditions used to culture recipient bacteria. 10
Supplementary Table 3. Conjugation colony counts. 10
Supplementary Table 4. Conjugation frequency data. 11
Supplementary Table 5. DNA extraction data for all organisms. 12
Supplementary Table 6. Detailed summary of ORI-marker screen results in all strains. 12
Supplementary Table 7. Colony counts from selective plates used for NGS. 12
Supplementary Table 8. Oligonucleotides used for amplicon sequencing. 13

### Supplementary Figures

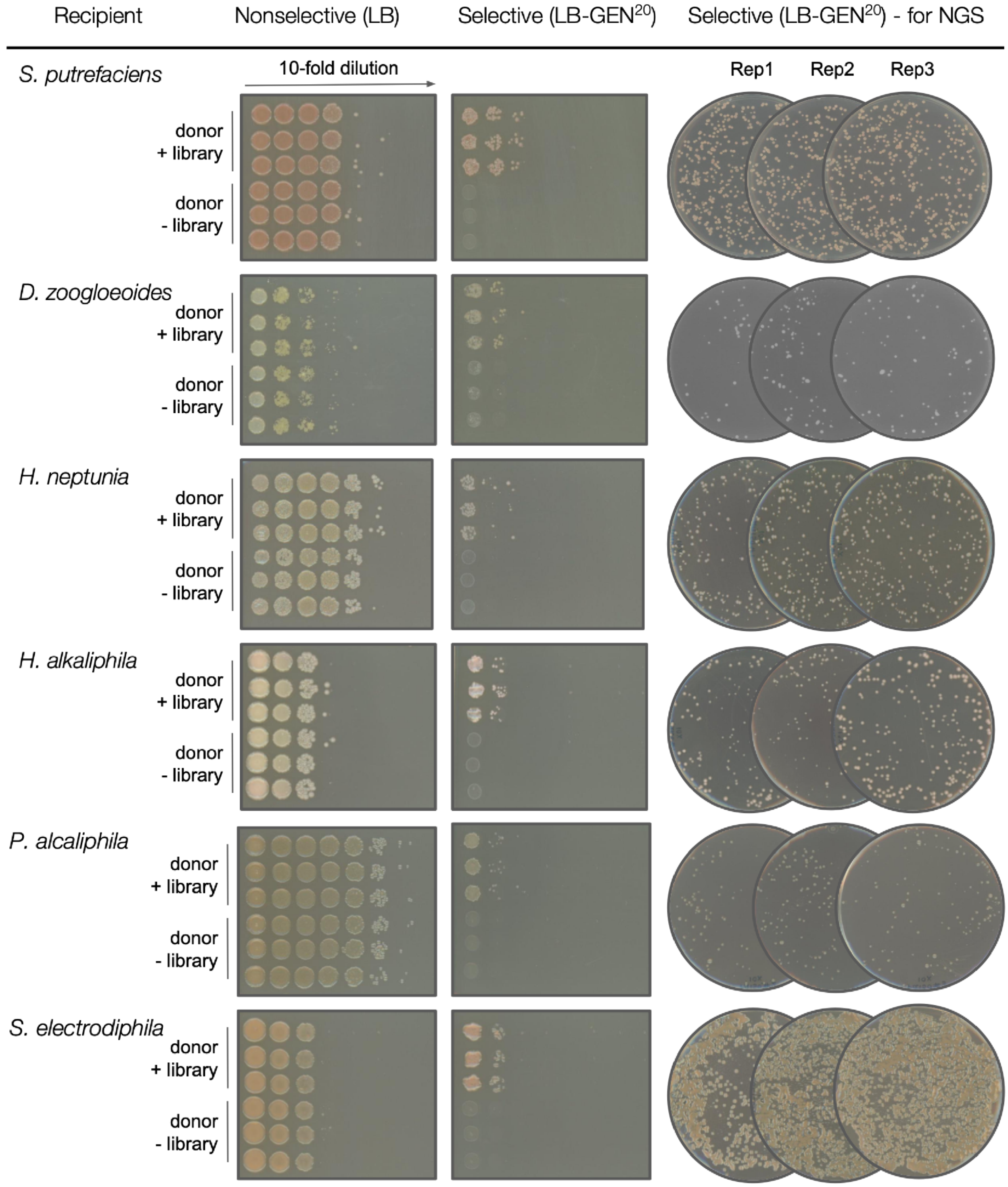

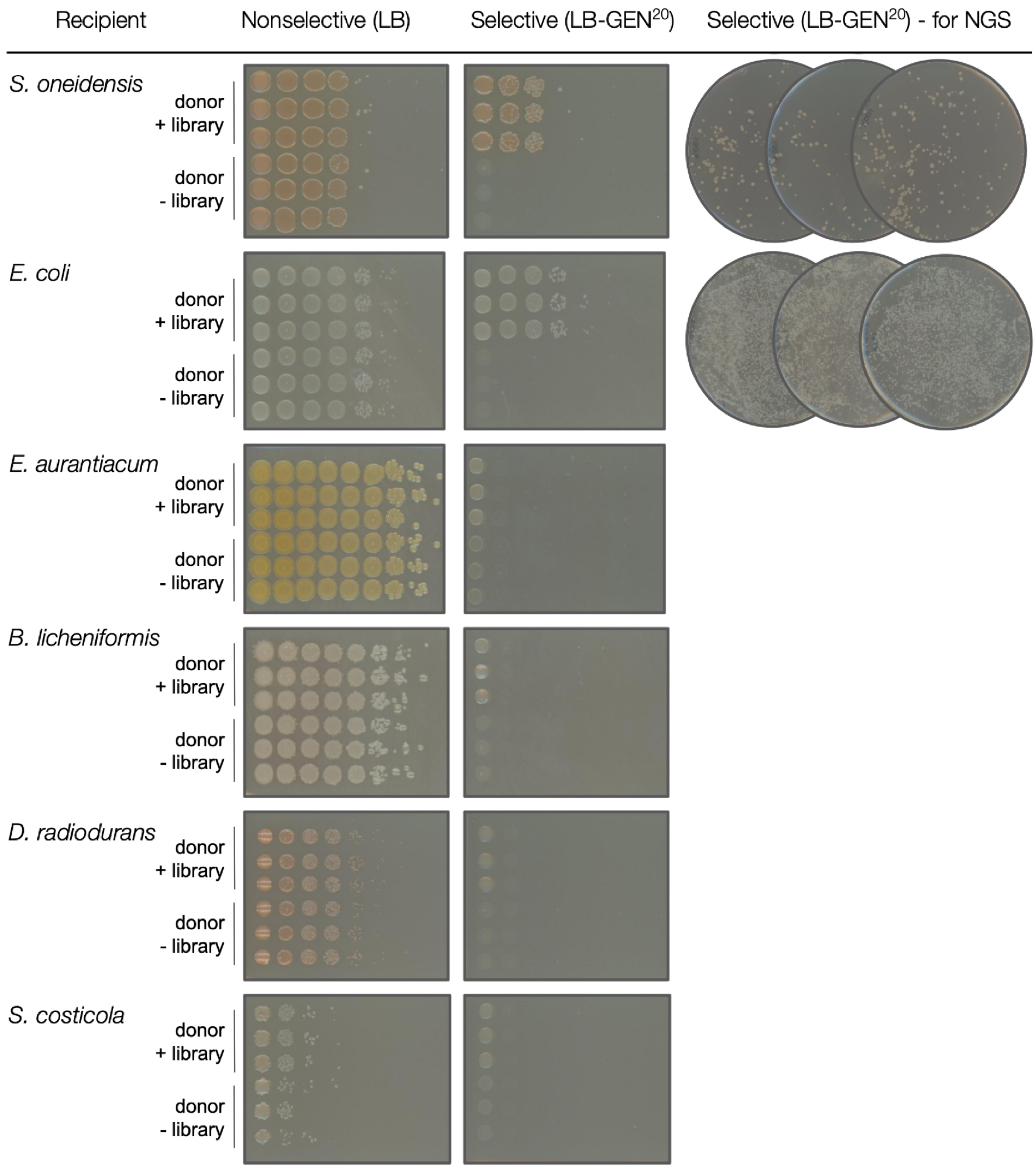


**Supplementary Figure 1. Conjugation plate images - Library pGL2_150.**
Images of spot plates and full plates for 12 bacteria following the ORI-marker screen with pGL2_150. Conjugations were performed in triplicate with an *E. coli* donor harboring the pGL2_150 library (donor + library) or without the library (donor - library).

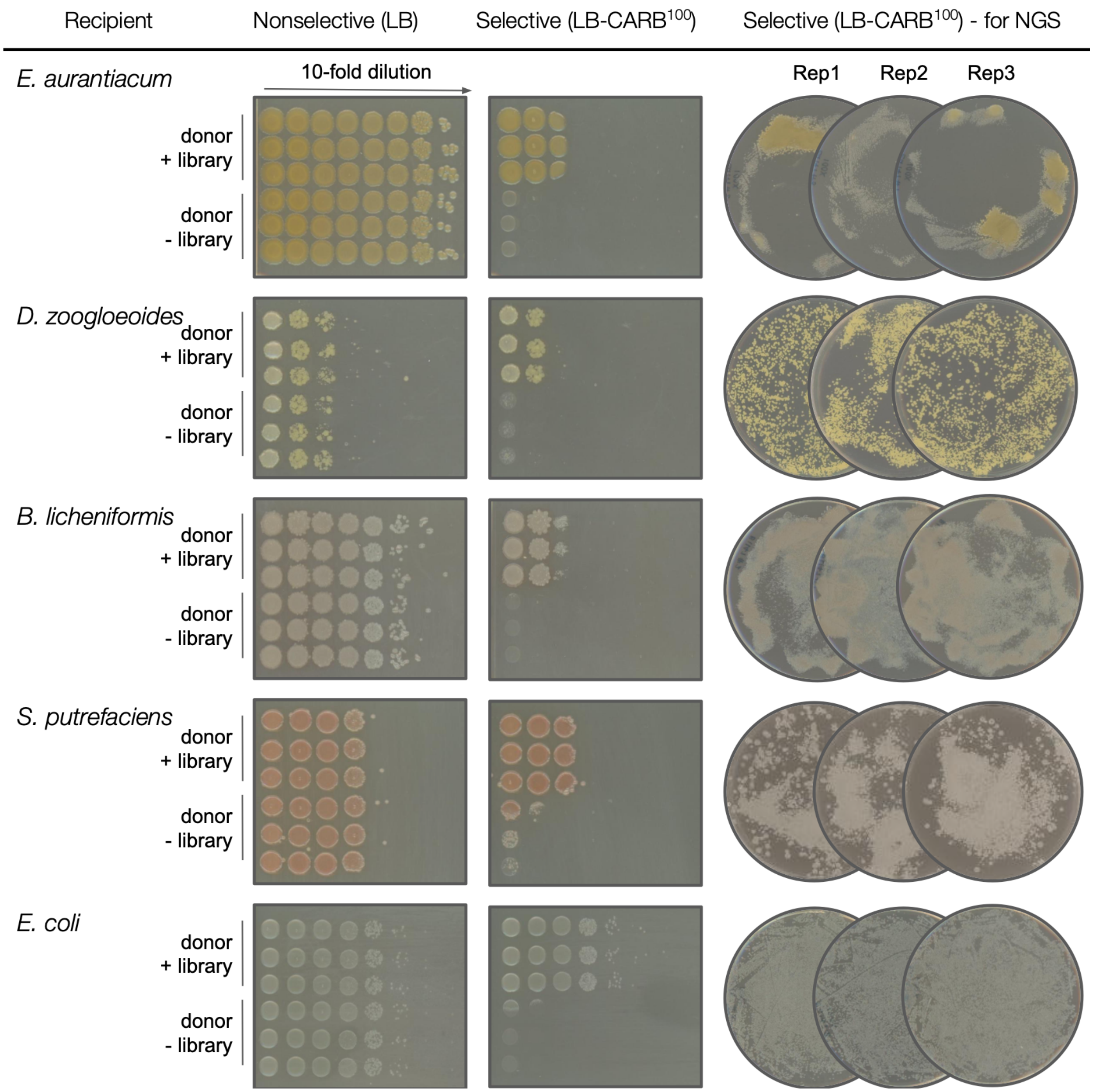


**Supplementary Figure 2. Conjugation plate images - Library pGL2_147**

Images of spot plates and full plates for 5 bacteria following the ORI-marker screen with pGL2_147. Conjugations were performed in triplicate with an *E. coli* donor harboring the pGL2_147 library (donor + library) or without the library (donor - library).

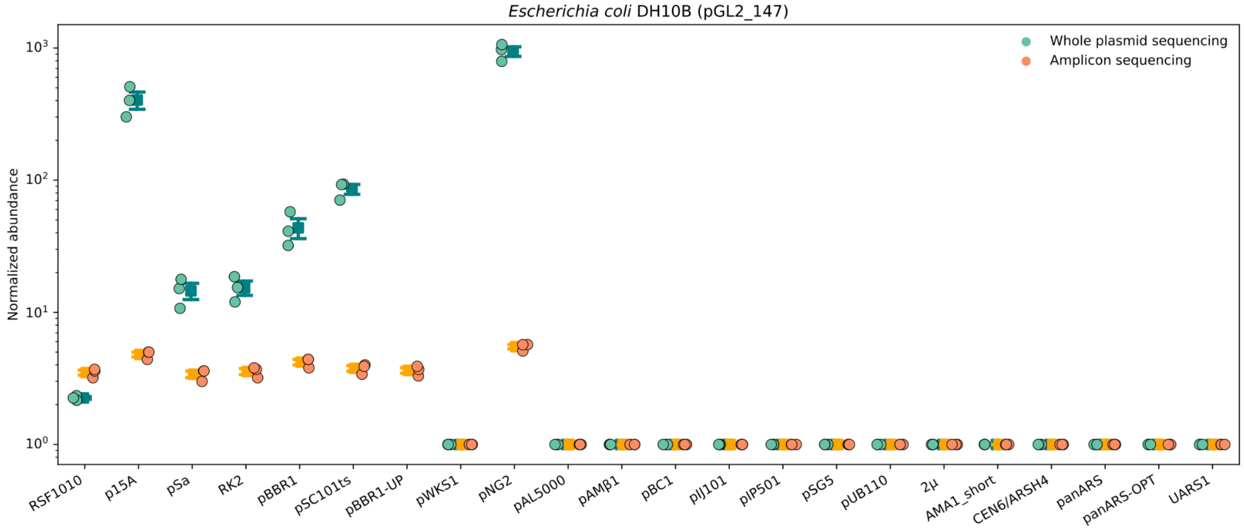
*Escherichia coli* DH10B (pGL2_147)
Whole plasmid sequencing
Amplicon sequencing
Normalized abundance
$10^0$
$10^1$
$10^2$
$10^3$
RSF1010
p15A
pSa
RK2
pBBR1
pSC101ts
pBBR1-UP
pWKS1
pNG2
pAL5000
pAMβ1
pBC1
pIJ101
pIP501
pSG5
pUB110
2μ
AMA1_short
CEN6/ARSH4
panARS
panARS-OPT
UARS1

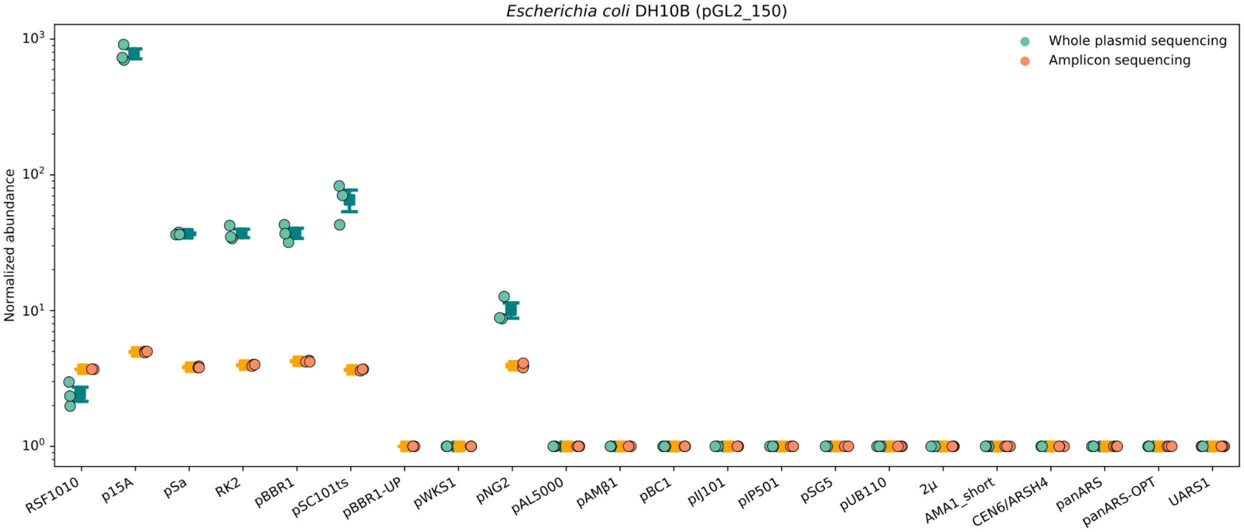
*Escherichia coli* DH10B (pGL2_150)
Whole plasmid sequencing
Amplicon sequencing
Normalized abundance
$10^0$
$10^1$
$10^2$
$10^3$
RSF1010
p15A
pSa
RK2
pBBR1
pSC101ts
pBBR1-UP
pWKS1
pNG2
pAL5000
pAMβ1
pBC1
pIJ101
pIP501
pSG5
pUB110
2μ
AMA1_short
CEN6/ARSH4
panARS
panARS-OPT
UARS1

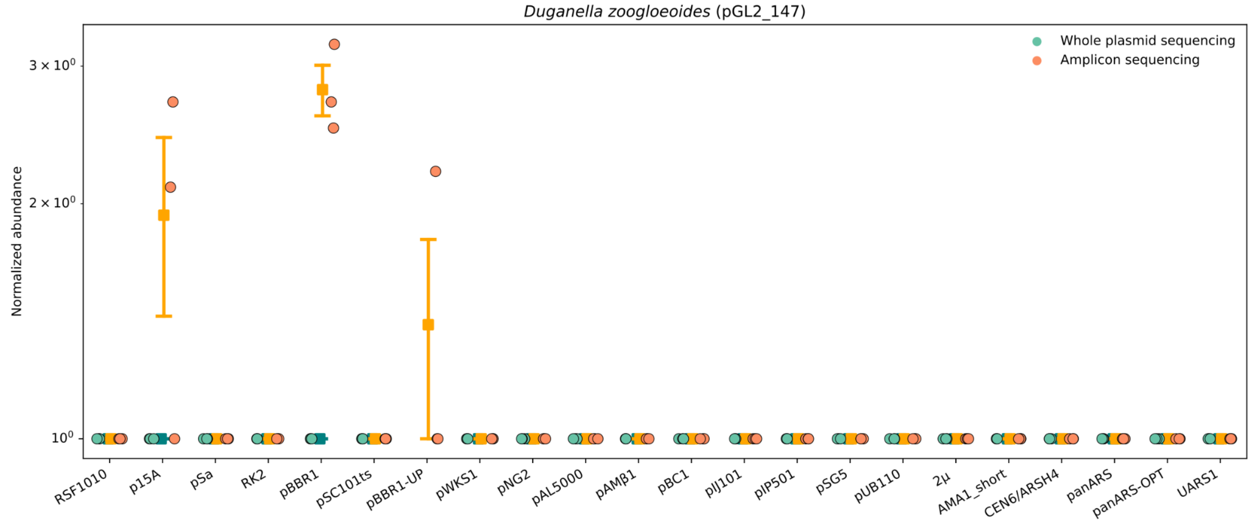
Duganella zoogloeoides (pGL2_147)
Whole plasmid sequencing
Amplicon sequencing
Normalized abundance
3 × 10⁰
2 × 10⁰
10⁰
RSF1010
p15A
pSa
RK2
pBBR1
pSC101ts
pBBR1-UP
pWKS1
pNG2
pAL5000
pAMβ1
pBC1
pIJ101
pIP501
pSG5
pUB110
2μ
AMA1_short
CEN6/ARSH4
panARS
panARS-OPT
UARS1

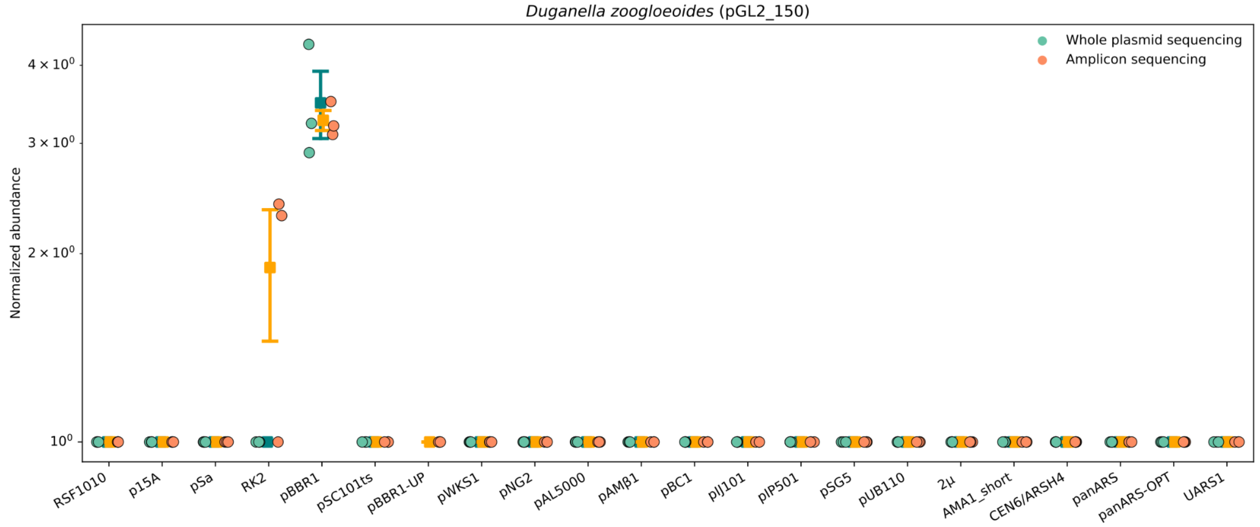
Duganella zoogloeoides (pGL2_150)
Whole plasmid sequencing
Amplicon sequencing
Normalized abundance
4 × 10⁰
3 × 10⁰
2 × 10⁰
10⁰
RSF1010
p15A
pSa
RK2
pBBR1
pSC101ts
pBBR1-UP
pWKS1
pNG2
pAL5000
pAMβ1
pBC1
pIJ101
pIP501
pSG5
pUB110
2μ
AMA1_short
CEN6/ARSH4
panARS
panARS-OPT
UARS1

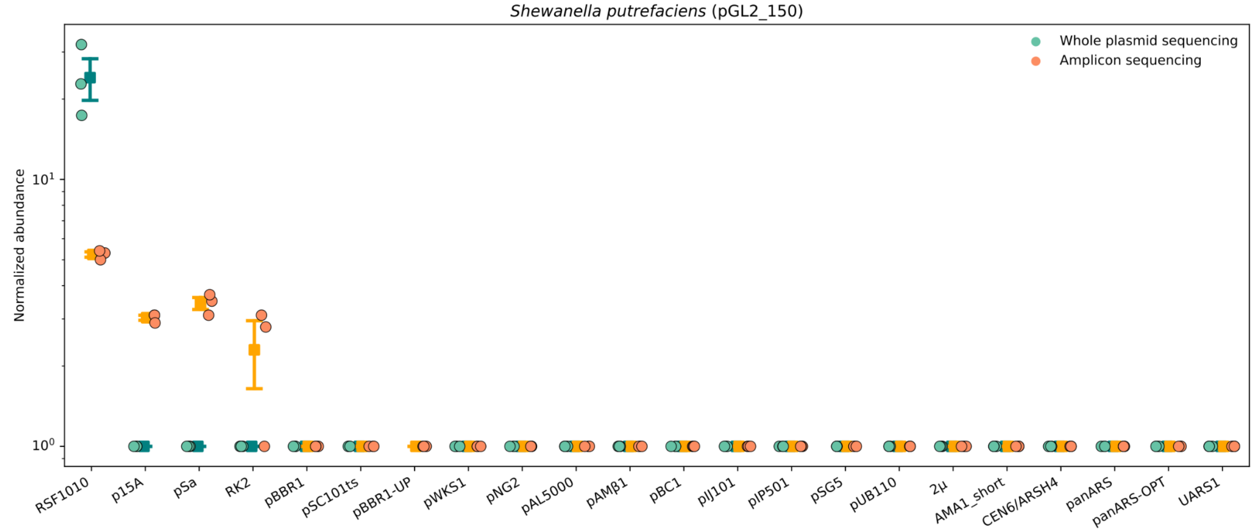
Shewanella putrefaciens (pGL2_150)
Whole plasmid sequencing
Amplicon sequencing
Normalized abundance
10¹
10⁰
RSF1010
p15A
pSa
RK2
pBBR1
pSC101ts
pBBR1-UP
pWKS1
pNG2
pAL5000
pAMβ1
pBC1
pIJ101
pIP501
pSG5
pUB110
2μ
AMA1_short
CEN6/ARSH4
panARS
panARS-OPT
UARS1

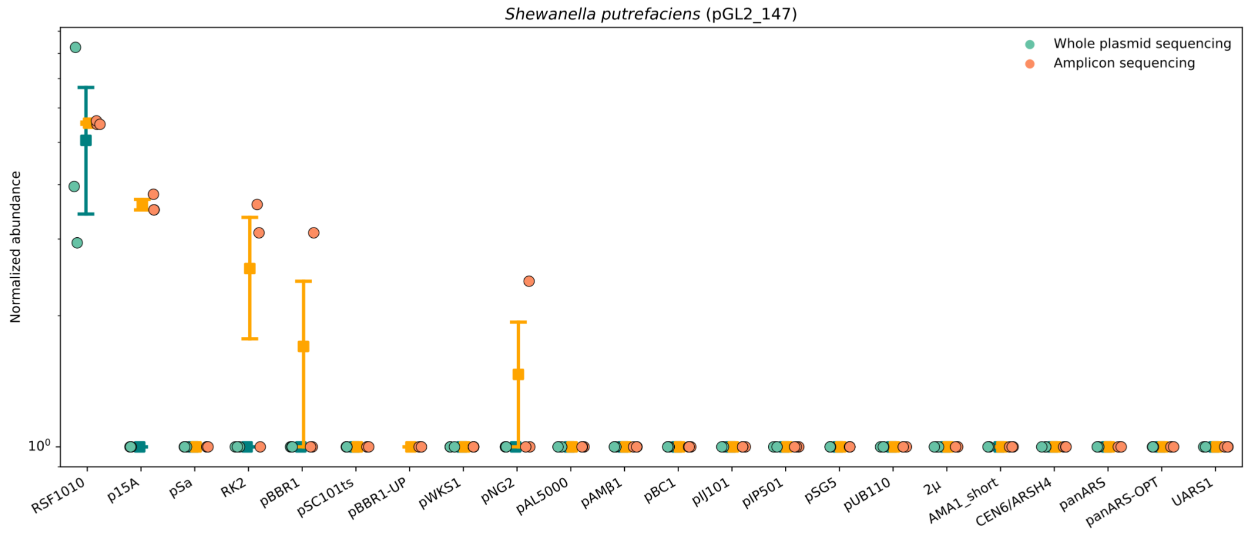
*Shewanella putrefaciens* (pGL2_147)
Whole plasmid sequencing
Amplicon sequencing
Normalized abundance
$10^0$
RSF1010
p15A
pSa
RK2
pBBR1
pSC101ts
pBBR1-UP
pWKS1
pNG2
pAL5000
pAMβ1
pBC1
pIJ101
pIP501
pSG5
pUB110
2μ
AMA1_short
CEN6/ARSH4
panARS
panARS-OPT
UARS1

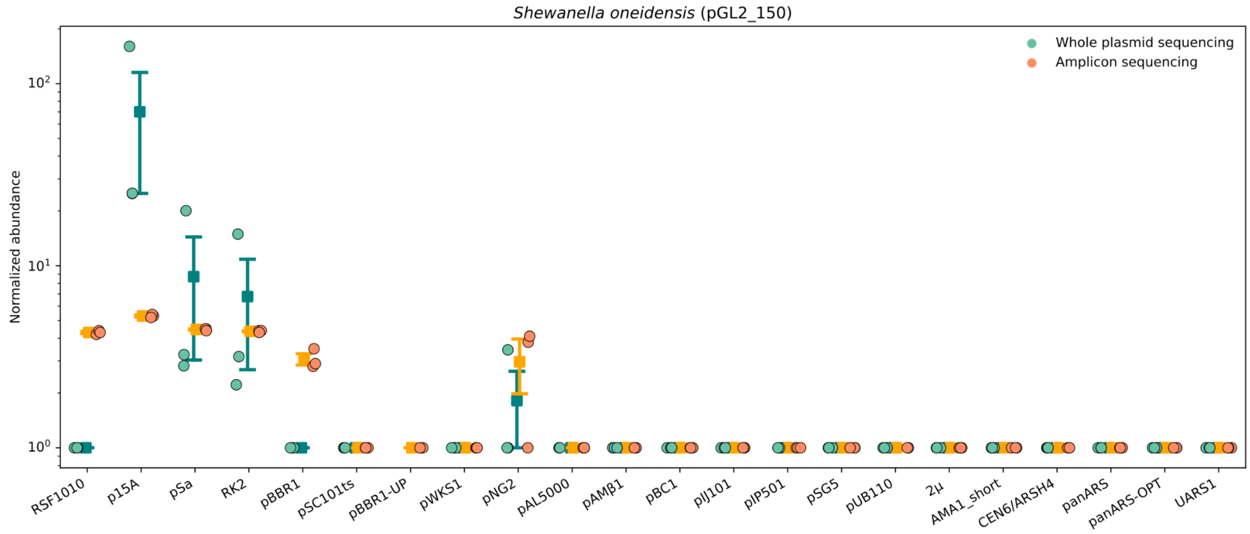
*Shewanella oneidensis* (pGL2_150)
Whole plasmid sequencing
Amplicon sequencing
Normalized abundance
$10^2$
$10^1$
$10^0$
RSF1010
p15A
pSa
RK2
pBBR1
pSC101ts
pBBR1-UP
pWKS1
pNG2
pAL5000
pAMβ1
pBC1
pIJ101
pIP501
pSG5
pUB110
2μ
AMA1_short
CEN6/ARSH4
panARS
panARS-OPT
UARS1

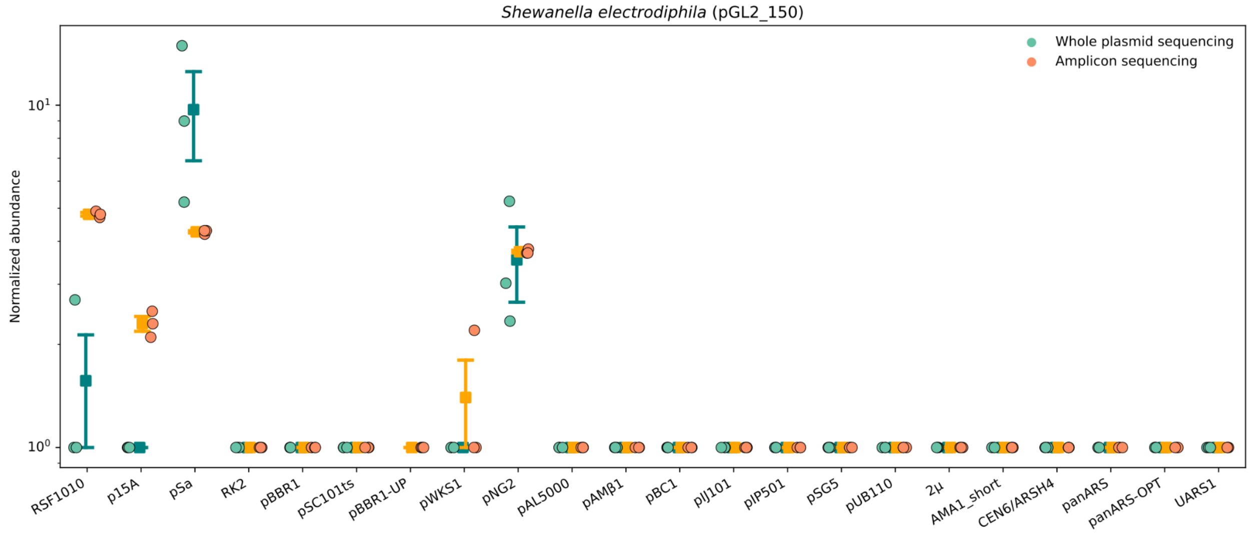
*Shewanella electrodiphila* (pGL2_150)
Whole plasmid sequencing
Amplicon sequencing
Normalized abundance
$10^1$
$10^0$
RSF1010
p15A
pSa
RK2
pBBR1
pSC101ts
pBBR1-UP
pWKS1
pNG2
pAL5000
pAMβ1
pBC1
pIJ101
pIP501
pSG5
pUB110
2μ
AMA1_short
CEN6/ARSH4
panARS
panARS-OPT
UARS1

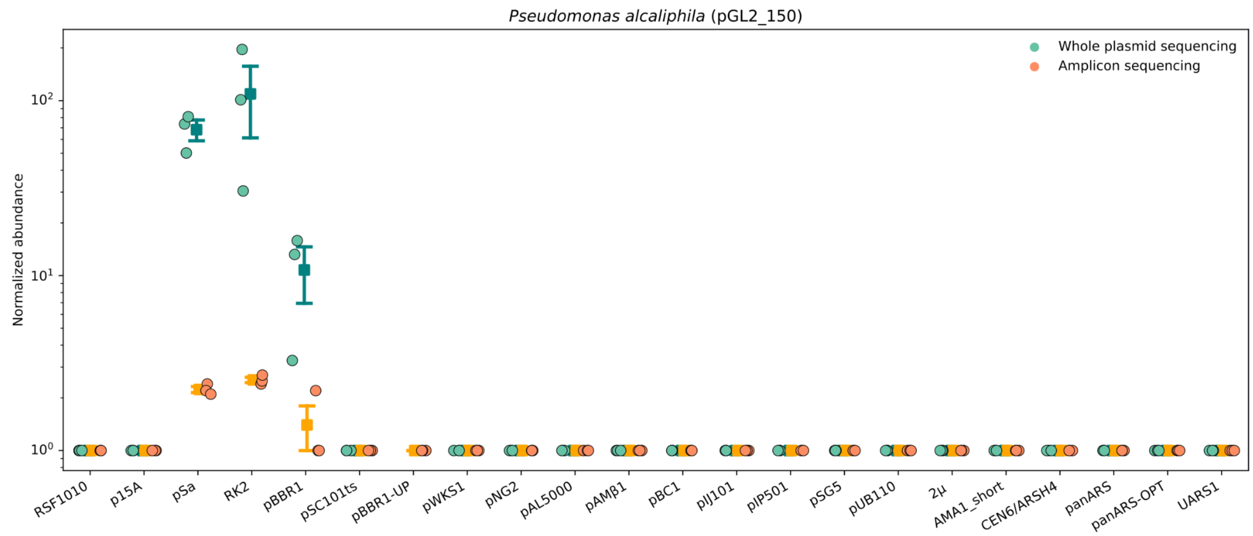
*Pseudomonas alcaliphila* (pGL2_150)
Whole plasmid sequencing
Amplicon sequencing
Normalized abundance
$10^2$
$10^1$
$10^0$
RSF1010
p15A
pSa
RK2
pBBR1
pSC101ts
pBBR1-UP
pWKS1
pNG2
pAL5000
pAMβ1
pBC1
pIJ101
pIP501
pSG5
pUB110
2μ
AMA1_short
CEN6/ARSH4
panARS
panARS-OPT
UARS1

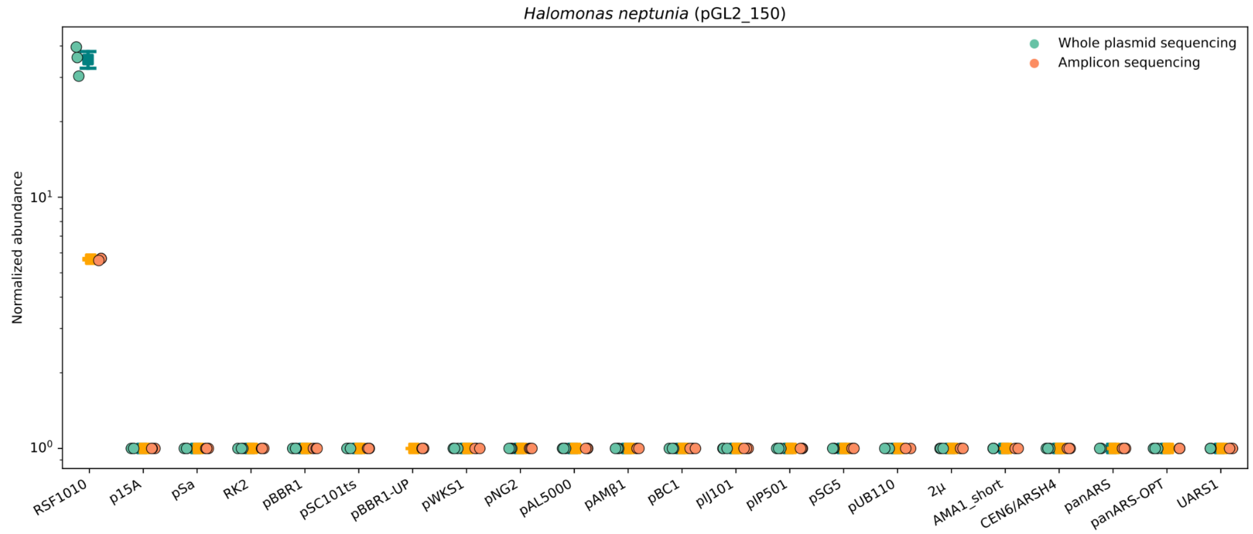
*Halomonas neptunia* (pGL2_150)
Whole plasmid sequencing
Amplicon sequencing
Normalized abundance
$10^1$
$10^0$
RSF1010
p15A
pSa
RK2
pBBR1
pSC101ts
pBBR1-UP
pWKS1
pNG2
pAL5000
pAMβ1
pBC1
pIJ101
pIP501
pSG5
pUB110
2μ
AMA1_short
CEN6/ARSH4
panARS
panARS-OPT
UARS1

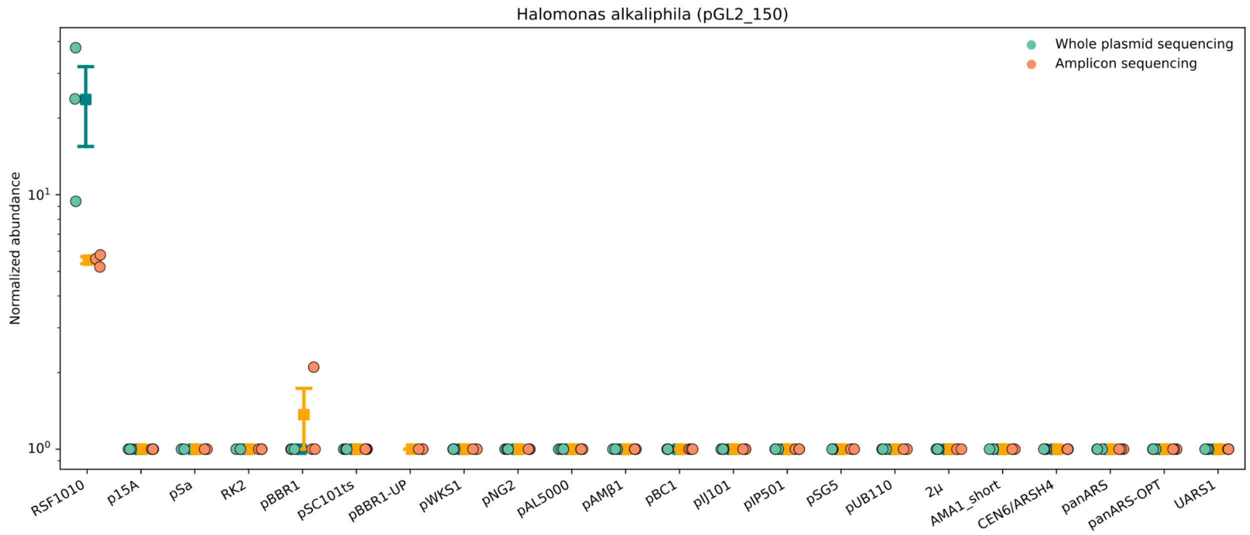
Halomonas alkaliphila (pGL2_150)
Whole plasmid sequencing
Amplicon sequencing
Normalized abundance
$10^1$
$10^0$
RSF1010
p15A
pSa
RK2
pBBR1
pSC101ts
pBBR1-UP
pWKS1
pNG2
pAL5000
pAMβ1
pBC1
pIJ101
pIP501
pSG5
pUB110
2μ
AMA1_short
CEN6/ARSH4
panARS
panARS-OPT
UARS1

**Supplementary Figure 3. Relative ORI abundances in each recipient bacteria.**
Results based on each sequencing method are indicated as whole plasmid sequencing (teal) or amplicon sequencing (orange). Each point represents a sequenced biological replicate for that strain (n=3). Amplicon results are normalized to the non-functional control ORI as described in *Methods*, and whole plasmid sequencing results are represented by the mean depth of coverage of each ORI. Error bars represent standard error of the mean.

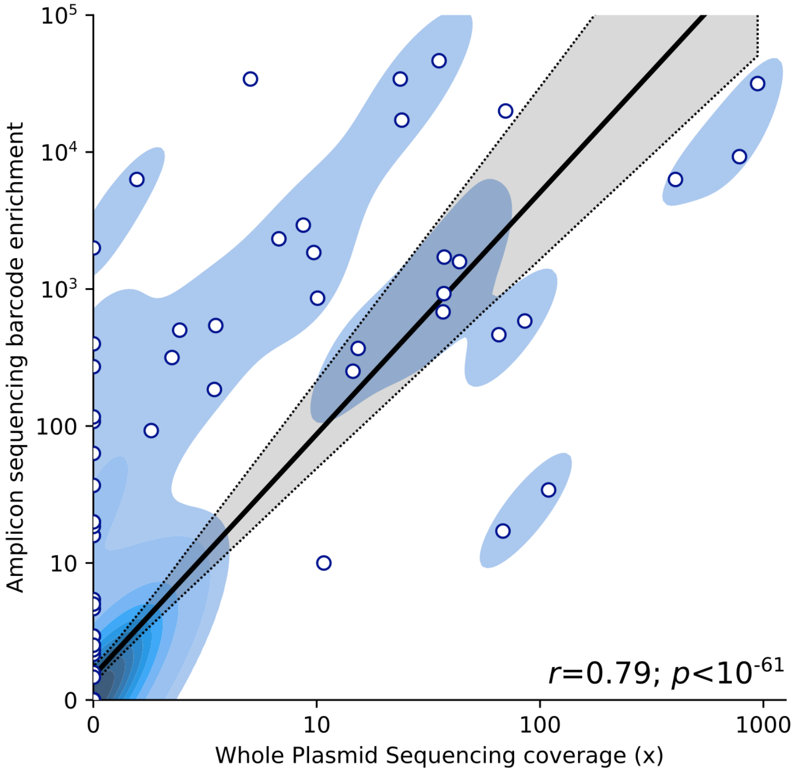


**Supplementary Figure 4. Correlation between amplicon and whole plasmid sequencing.**
Correlation of amplicon sequencing with whole plasmid sequencing for ORI detection. Each point represents the abundance of a given origin from a transconjugant pool as sequenced both through the amplicon approach and whole plasmid sequencing. n=39 samples from n=10 different species are included in the plot.

### Supplementary Tables

**Supplementary Table 1. Nucleotide sequences of the ORIs used in this study.**
This table is available as an .xls file accompanying the online manuscript.

**Supplementary Table 2. Conditions used to culture recipient bacteria.**
All organisms were grown in LB media in liquid or agar plates prior to conjugation as indicated.

| CVM ID | Strain | Culture method | Growth temperature (°C) |
|---|---|---|---|
| CVM007 | *Shewanella oneidensis* | liquid | 30 |
| CVM010 | *Pseudomonas alcaliphila* | liquid | 30 |
| CVM021 | *Deinococcus radiodurans* | liquid | 30 |
| CVM022 | *Bacillus licheniformis* | liquid | 30 |
| CVM027 | *Halomonas neptunia* | agar | 30 |
| CVM028 | *Exiguobacterium aurantiacum* | liquid | 37 |
| CVM029 | *Salinivibrio costicola subsp. alcaliphilus* | agar | 30 |
| CVM032 | *Halomonas alkaliphila* | agar | 37 |
| CVM036 | *Shewanella electrodiphila* | agar | 15 |
| CVM037 | *Shewanella putrefaciens* | liquid | 30 |
| CVM038 | *Duganella zoogloeoides* | liquid | 30 |
| CVM080 | *Escherichia coli* | liquid | 37 |

**Supplementary Table 3. Conjugation colony counts.**
Colony counts observed in the ORI-marker screen in 12 bacteria. Counts are provided in triplicate (Rep 1-3) for the indicated library, for both selective and nonselective spot plates.

| CVM ID | Organism | ORI-marker library | Spot plate colony counts | | | | | |
|---|---|---|---|---|---|---|---|---|
| | | | Rep 1 | | Rep 2 | | Rep 3 | |
| | | | Selective | Nonselective | Selective | Nonselective | Selective | Nonselective |
| CVM007 | *S. oneidensis* | pGL2_150 | $35 \times 10^2$ | $3 \times 10^4$ | $32 \times 10^2$ | $5 \times 10^4$ | $25 \times 10^2$ | $2 \times 10^4$ |
| CVM010 | *P. alcaliphila* | pGL2_150 | $10 \times 10^1$ | $3 \times 10^6$ | $9 \times 10^1$ | $2 \times 10^6$ | $7 \times 10^1$ | $2 \times 10^6$ |
| CVM021 | *D. radiodurans* | pGL2_150 | 0 | $6 \times 10^5$ | 0 | $4 \times 10^5$ | 0 | $6 \times 10^5$ |
| CVM022 | *B. licheniformis* | pGL2_147 | $23 \times 10^2$ | $15 \times 10^5$ | $17 \times 10^2$ | $7 \times 10^5$ | $8 \times 10^2$ | $7 \times 10^5$ |
| | | pGL2_150 | 0 | $10 \times 10^6$ | 0 | $10 \times 10^6$ | 0 | $5 \times 10^6$ |
| CVM027 | *H. neptunia* | pGL2_150 | $5 \times 10^1$ | $7 \times 10^2$ | $3 \times 10^1$ | $2 \times 10^2$ | $2 \times 10^1$ | $4 \times 10^2$ |
| CVM028 | *E. aurantiacum* | pGL2_147 | *$10 \times 10^2$ | $4 \times 10^7$ | *$10 \times 10^2$ | $5 \times 10^7$ | *$10 \times 10^2$ | $7 \times 10^7$ |
| | | pGL2_150 | 0 | $7 \times 10^7$ | 0 | $8 \times 10^7$ | 0 | $1 \times 10^7$ |
| CVM029 | *S. costicola subsp. alkaliphilus* | pGL2_150 | 0 | $2 \times 10^3$ | 0 | $1 \times 10^3$ | 0 | $1 \times 10^3$ |

| CVM ID | Organism | ORI-marker library | Spot plate colony counts | | | | | |
|---|---|---|---|---|---|---|---|---|
| | | | Rep 1 | | Rep 2 | | Rep 3 | |
| | | | Selective | Nonselective | Selective | Nonselective | Selective | Nonselective |
| CVM032 | *H. alkaliphila* | pGL2_150 | 7 x $10^1$ | 51 x $10^2$ | 17 x $10^1$ | 24 x $10^2$ | 9 x $10^1$ | 32 x $10^2$ |
| CVM036 | *S. electrodiphila* | pGL2_150 | 15 x $10^1$ | *1 x $10^3$ | 14 x $10^1$ | 2 x $10^3$ | 15 x $10^1$ | *1 x $10^3$ |
| CVM037 | *S. putrefaciens* | pGL2_147 | *1 x $10^3$ | 1 x $10^4$ | *1 x $10^3$ | *1 x $10^4$ | 1 x $10^3$ | 1 x $10^4$ |
| | | pGL2_150 | 6 x $10^2$ | 1 x $10^4$ | 8 x $10^2$ | 1 x $10^4$ | 5 x $10^2$ | 3 x $10^4$ |
| CVM038 | *D. zoogloeoides* | pGL2_147 | 52 x $10^1$ | 4 x $10^3$ | 47 x $10^1$ | 5 x $10^3$ | 54 x $10^1$ | 5 x $10^3$ |
| | | pGL2_150 | 11 x $10^1$ | 6 x $10^3$ | 8 x $10^1$ | 5 x $10^3$ | 15 x $10^1$ | 10 x $10^3$ |
| CVM080 | *E. coli DH10B* | pGL2_147 | 3 x $10^5$ | 9 x $10^5$ | 3 x $10^5$ | 18 x $10^5$ | 1 x $10^5$ | 8 x $10^5$ |
| | | pGL2_150 | 1 x $10^4$ | 8 x $10^5$ | 4 x $10^4$ | 11 x $10^5$ | 3 x $10^4$ | 4 x $10^5$ |

*Colonies were too merged to count, therefore a single colony was reported for one dilution higher than the dilution where growth was observed.

**Supplementary Table 4. Conjugation frequency data.**

Conjugation frequency reported for three replicates (Rep 1-3) of the ORI-marker screen in 12 bacteria with the pGL2_147 and/or pGL2_150 ORI-marker library. SD, standard deviation.

| CVM ID | Organism | ORI-marker library | Conjugation Frequency (transconjugants/recipient) | | | | |
|---|---|---|---|---|---|---|---|
| | | | Rep 1 | Rep 2 | Rep 3 | Average | SD |
| CVM007 | *S. oneidensis* | pGL2_150 | 0.117 | 0.064 | 0.125 | 0.102 | 0.033 |
| CVM010 | *P. alcaliphila* | pGL2_150 | 0.000033 | 0.000045 | 0.000035 | 0.000038 | 0.000006 |
| CVM021 | *D. radiodurans* | pGL2_150 | 0 | 0 | 0 | 0 | 0 |
| CVM022 | *B. licheniformis* | pGL2_147 | 0.00153 | 0.00243 | 0.00114 | 0.00170 | 0.00066 |
| | | pGL2_150 | 0 | 0 | 0 | 0 | 0 |
| CVM027 | *H. neptunia* | pGL2_150 | 0.0714 | 0.1500 | 0.0500 | 0.0905 | 0.0527 |
| CVM028 | *E. aurantiacum* | pGL2_147 | 0.000025 | 0.00002 | 0.000014 | 0.000020 | 0.00001 |
| | | pGL2_150 | 0 | 0 | 0 | 0 | 0 |
| CVM029 | *S. costicola subsp. alkaliphilus* | pGL2_150 | 0 | 0 | 0 | 0 | 0 |
| CVM032 | *H. alkaliphila* | pGL2_150 | 0.0137 | 0.0708 | 0.0281 | 0.0376 | 0.0297 |
| CVM036 | *S. electrodiphila* | pGL2_150 | 0.150 | 0.070 | 0.150 | 0.123 | 0.046 |
| CVM037 | *S. putrefaciens* | pGL2_147 | 0.1 | 0.1 | 0.1 | 0.1 | 0.0 |
| | | pGL2_150 | 0.060 | 0.080 | 0.020 | 0.053 | 0.031 |
| CVM038 | *D. zoogloeoides* | pGL2_147 | 0.130 | 0.090 | 0.110 | 0.110 | 0.020 |
| | | pGL2_150 | 0.02 | 0.02 | 0.02 | 0.02 | 0.00 |
| CVM080 | *E. coli* | pGL2_147 | 0.333 | 0.167 | 0.125 | 0.208 | 0.110 |
| | | pGL2_150 | 0.0125 | 0.0364 | 0.0750 | 0.0413 | 0.0315 |

**Supplementary Table 5. DNA extraction data for all organisms.**
DNA concentrations from plasmid and genomic DNA extractions for 12 bacteria following the ORI-marker screen with pGL2_147 and/or pGL2_150. SD, standard deviation.

| Organism | Library | Plasmid (ng/µL) | | | | | gDNA (ng/µL) | | | | |
|---|---|---|---|---|---|---|---|---|---|---|---|
| | | Rep 1 | Rep 2 | Rep 3 | Avg | SD | Rep 1 | Rep 2 | Rep 3 | Avg | SD |
| *S. oneidensis* | pGL2_150 | 77.3 | 173.0 | 181.0 | 143.8 | 57.7 | 309.0 | 225.0 | 309.0 | 281.0 | 48.5 |
| *P. alcaliphila* | pGL2_150 | 5.2 | 2.8 | 4.6 | 4.2 | 1.2 | 335.0 | 303.0 | 331.0 | 323.0 | 17.4 |
| *B. licheniformis* | pGL2_147 | 23.5 | 26.3 | 18.5 | 22.8 | 4.0 | 290.0 | 141.0* | 129.0 | 186.7 | 89.7 |
| *H. neptunia* | pGL2_150 | 6.9 | 8.7 | 8.3 | 8.0 | 0.9 | 60.1 | 156.0 | 31.1 | 82.4 | 65.4 |
| *E. aurantiacum* | pGL2_147 | 11.2 | 7.2 | 3.8 | 7.4 | 3.7 | 654.0 | 673.0* | 385.0 | 570.7 | 161.1 |
| *H. alkaliphila* | pGL2_150 | 8.5 | 8.1 | 4.0 | 6.8 | 2.5 | 249.0 | 88.7 | 216.0 | 184.6 | 84.6 |
| *S. electrodiphila* | pGL2_150 | 6.5 | 10.1 | 9.5 | 8.7 | 2.0 | 376.0* | 385.0* | 373.0* | 378.0 | 6.2 |
| *S. putrefaciens* | pGL2_147 | 189.0 | 161.0 | 161.0 | 170.3 | 16.2 | 11.3 | 20.2 | 5.6 | 12.4 | 7.4 |
| | pGL2_150 | 153.0 | 153.0 | 164.0 | 156.7 | 6.4 | 37.5 | 49.5 | 60.1 | 49.0 | 11.3 |
| *D. zoogloeoides* | pGL2_147 | 86.7 | 123.0 | 63.5 | 91.1 | 30.0 | 189.0* | 378.0* | 96.0* | 221.0 | 143.7 |
| | pGL2_150 | 69.3 | 46.9 | 66.7 | 61.0 | 12.3 | 693.0* | 767.0 | 425.0* | 628.3 | 179.9 |
| *E. coli* | pGL2_147 | 68.0 | 73.3 | 54.5 | 65.3 | 9.7 | 178.0 | 116.0 | 163.0 | 152.3 | 32.3 |
| | pGL2_150 | 41.7 | 34.7 | 34.8 | 37.1 | 4.0 | 44.1 | 87.3 | 142.0 | 91.1 | 49.1 |
| Control ($10^{-1}$ dilution) | pGL2_150 | 0.0767 | 0.151 | 0.157 | 0.1 | 0.0 | 2.89 | 9.67 | 4.71 | 5.8 | 3.5 |
| Control ($10^{-2}$ dilution) | pGL2_150 | 0.00000973 | 0.000465 | 0.0853 | 0.0286 | 0.0491 | 0.331 | 0.212 | 0.209 | 0.251 | 0.070 |

*Poor magnetic bead separation during the elution step of gDNA extraction, therefore an additional 200 µL of elution buffer was added.

**Supplementary Table 6. Detailed summary of ORI-marker screen results in all strains.**
This table is available as an .xls file accompanying the online manuscript.

**Supplementary Table 7. Colony counts from selective plates used for NGS.**
Colony counts observed on full selective plates following the ORI-marker screen in 12 bacteria. Counts are provided in triplicate (Rep 1-3) for the indicated library. For plates where colonies were too numerous to count 1000+ colonies were reported. SD, standard deviation.

| CVM ID | Organism | ORI-marker library | Full plate colony counts | | | | |
|---|---|---|---|---|---|---|---|
| | | | Rep 1 | Rep 2 | Rep 3 | Average | SD |
| CVM007 | *Shewanella oneidensis* | pGL2_150 | 90 | 222 | 230 | 180 | 78.6 |
| CVM010 | *Pseudomonas alcaliphila* | pGL2_150 | 156 | 93 | 126 | 125 | 31.5 |
| CVM022 | *Bacillus licheniformis* | pGL2_147 | 1000+ | 1000+ | 1000+ | 1000+ | n/a |
| CVM027 | *Halomonas neptunia* | pGL2_150 | 386 | 350 | 278 | 338 | 55 |

| CVM ID | Organism | ORI-marker library | Full plate colony counts | | | | |
|---|---|---|---|---|---|---|---|
| | | | Rep 1 | Rep 2 | Rep 3 | Average | SD |
| CVM028 | *Exiguobacterium aurantiacum* | pGL2_147 | 1000+ | 1000+ | 1000+ | 1000+ | n/a |
| CVM032 | *Halomonas alkaliphila* | pGL2_150 | 171 | 242 | 147 | 187 | 49.4 |
| CVM036 | *Shewanella electrodiphila* | pGL2_150 | 1000+ | 1000+ | 1000+ | 1000+ | n/a |
| CVM037 | *Shewanella putrefaciens* | pGL2_147 | 1000+ | 1000+ | 1000+ | 1000+ | n/a |
| | | pGL2_150 | 549 | 476 | 540 | 521 | 40 |
| CVM038 | *Duganella zoogloeoides* | pGL2_147 | 1000+ | 1000+ | 1000+ | 1000+ | n/a |
| | | pGL2_150 | 110 | 60 | 102 | 91 | 27 |
| CVM080 | *E. coli* | pGL2_147 | 1000+ | 1000+ | 1000+ | 1000+ | n/a |
| | | pGL2_150 | 1000+ | 1000+ | 1000+ | 1000+ | n/a |

**Supplementary Table 8. Oligonucleotides used for amplicon sequencing.**
Primers used for "PCR 1" amplification as part of the amplicon sequencing workflow. These primers bind to constant regions upstream and downstream of the unique barcode sequences associated with each plasmid ORI. The underlined regions represent 5' primer extensions which introduce stub sequences enabling binding of dual indexed Illumina i5 and i7 primers in the subsequent "PCR 2" step.

| Primer | Sequence | Purpose |
|---|---|---|
| oCG0038 | TCCCTACACGACGCTCTTCCGATCTGGGTCACGCGTAGGACG | PCR 1 forward |
| oCG0039 | GAGTTCAGACGTGTGCTCTTCCGATCTCCAGCTTCACACGGCGT | PCR 1 reverse |
